# CTCF-anchored chromatin loop dynamics during human meiosis

**DOI:** 10.1101/2024.02.29.582729

**Authors:** Vera B. Kaiser, Colin A. Semple

## Abstract

**Background:** During meiosis, the mammalian genome is organised within chromatin loops, which facilitate synapsis, crossing over and chromosome segregation, setting the stage for recombination events and the generation of genetic diversity. Chromatin looping is thought to play a major role in the establishment of cross overs during prophase I of meiosis, in diploid early primary spermatocytes. However, chromatin conformation dynamics during human meiosis are difficult to study experimentally, due to the transience of each cell division and the difficulty of obtaining stage-resolved cell populations. Here, we employed a machine learning framework trained on single cell ATAC-seq and RNA-seq data to predict CTCF-anchored looping during spermatogenesis, including cell types at different stages of meiosis.

**Results:** We find dramatic changes in genome-wide looping patterns throughout meiosis: compared to pre-and-post meiotic germline cell types, loops in meiotic early primary spermatocytes are more abundant, more variable between individual cells, and more evenly spread throughout the genome. In preparation for the first meiotic division, loops also include longer stretches of DNA, encompassing more than half of the total genome. These loop structures then influence the rate of recombination initiation and resolution as cross overs. In contrast, in later mature sperm stages, we find evidence of genome compaction, with loops being confined to the telomeric ends of the chromosomes.

**Conclusion:** Overall, we find that chromatin loops do not orchestrate the gene expression dynamics seen during spermatogenesis, but loops do play important roles in recombination, influencing the positions of DNA breakage and cross over events.

## BACKGROUND

Recombination, the generation of novel combinations of alleles, is a critical process that affects an individual organism’s phenotype, population-level amounts of genetic diversity and the response to selection (1, 2). In mammals, homologous recombination occurs during meiosis, the cytological process that gives rise to gametes which provide all genetic information for the next generation (3, 4). Prophase I of meiosis is a lengthy and complex process and is subdivided into five different stages (leptotene, zygotene, pachytene, diplotene, and diakinesis), which, together, last for several days in human males. Prophase I takes place in primary spermatocytes, where programmed DNA double-strand breaks (DSBs) are introduced, homologous chromosomes pair at the synaptonemal complex and exchange genetic material (5). When DSBs occur during pachytene, overhanging single-stranded DNA (ssDNA) is bound by DMC1 near the breakage site (6). A fraction of such DMC1-bound ssDNA sites result in strand exchange and crossovers (7), but the underlying process by which sites are chosen remains unclear (8). At the population level, cross-over events are concentrated in recombination hotspots (HSs), which are typically 1-2kb in size (9–11), suggesting strong biases. Previously (12), we showed that HSs may be a by-product of particular chromatin environments, defined by patterns of chromatin looping, but chromatin data from human meiotic cells were not available at the time. However, the recent emergence of scATAC-seq data from human testicular samples (13) has provided new opportunities to study the dependencies between the process of recombination and chromatin structure in humans.

Chromatin looping is a fundamental level of organisation in all eukaryotic cells, including germline cells. In somatic cells, chromatin topology can be studied by methods such as Hi-C (14), ChIA-PET (15) and related methods. In the current widely accepted model, the cohesin complex catalyses the formation of chromatin loops at the scale of hundreds of kilobases (kb), and loops are anchored at convergently oriented CTCF-binding sites, which prevent the cohesin molecules from releasing the DNA (16). During interphase, chromatin loops appear to regulate gene expression (17); during meiosis, they may play a crucial role in a different context - the pairing of the homologous chromosomes at the synaptonemal complex. In mice, stage-resolved Hi-C analysis of spermatogenesis has been performed via the experimental synchronization of germ cells (18, 19); this has demonstrated that, while larger chromatin domains are lost in prophase I (20, 21), CTCF retains its insulator function (22) and may be recruited to the synaptonemal complex (SC) via co-localization with RAD21/SMC1/ SMC3 (23–25), although the details of interaction between CTCF and the SC remain to be demonstrated. Chromatin loops are known to persist throughout meiosis (5), and their anchor points coincide with CTCF binding (26); however, the size of loops throughout prophase I is contentious (27) as loops sizes of 0.8 – 2Mb have been measured by Hi-C experiments in mice (18, 21, 28), but computational analyses (8) have revealed shorter loops in A compartments – similar to typical interphase loops, which are of the order of 200kb (29).

Overall, the topological organisation of the human genome during meiosis remains elusive. In particular, the positions of the genome that form loops during human meiosis are currently unknown, as is the level of variation in looping patterns between individual cells. In addition, the relationship between the chromatin structure of meiotic DNA and features of recombination (8) is largely unexplored.

Computational methods that employ machine learning strategies to predict chromatin looping (30–33) are a timely substitute when experimental data are hard to come by. Machine learning models can be trained on experimental data, such as CTCF-binding, gene expression and other epigenetic marks in combination with a set of “true” loops derived from Hi-C or ChiA-Pet datasets. Such models can predict chromatin looping with high accuracy – with AUC values typically > 95% - suggesting that computational approaches can sometimes outperform experimental data at predicting chromatin architecture, especially when sequencing depth is a limiting factor (31). However, in order to study chromatin looping during spermatogenesis, there is need to specifically study cells that are in the correct stage of meiosis, e.g. cells which undergo DSBs when studying the impact of looping on the process of recombination. One way to obtain sub-populations of cells is by single cell sequencing and selecting cells based on marker gene expression.

Here, we use machine learning methods to study a fundamental level of chromatin organisation during human meiosis: CTCF-anchored loops. We predict chromatin folding during all relevant cell stages and study the impact of chromatin loops on gene expression, DSB initiation, and homologous recombination patterns across the genome.

## RESULTS

### Dynamics of CTCF mediated chromatin loops during spermatogenesis

We interrogated a recently published testicular scATAC-seq dataset (13) to identify human meiotic cells. The cellranger-atac pipeline detected a combined total of 20,296 cells in the dataset (10,509 cells in SRR21861961, 7,086 in SRR21861962 and 2,701 in SRR21861963). Next, cell states were inferred using matched single cell RNA-seq data (34). Using marker gene expression, chromatin accessibility and the computation of “transfer anchors”, scRNA-seq profiles were used to project cell identities from the annotated scRNA data onto the scATAC-seq dataset (Fig. 1A) (35). As in Guo et al. (34), we classified cells into eight cell types of germ cell development: Spermatogonial Stem Cells (SSCs), Differentiating Spermatogonia, Early Primary Spermatocytes, Late Primary Spermatocytes, Round Spermatids, Elongated Spermatids, Sperm I and Sperm II. For each cell type, we identified between ∼ 700 to 4 thousand cells, including 2,008 early primary spermatocytes, i.e. diploid cells preparing for the first meiotic division. A pseudo time-course of meiosis is shown in the batch corrected UMAP plot (Fig. 1B) (36), displaying the transitions in chromatin accessibility across cells from SSCs to mature sperm. For illustrative purposes, we also show the scRNA-seq and scATAC-seq data projected onto the same plot (Fig. 1C and D), illustrating how cell identities were transferred from the gene expression dataset onto the chromatin accessibility dataset.

**Fig. 1:**
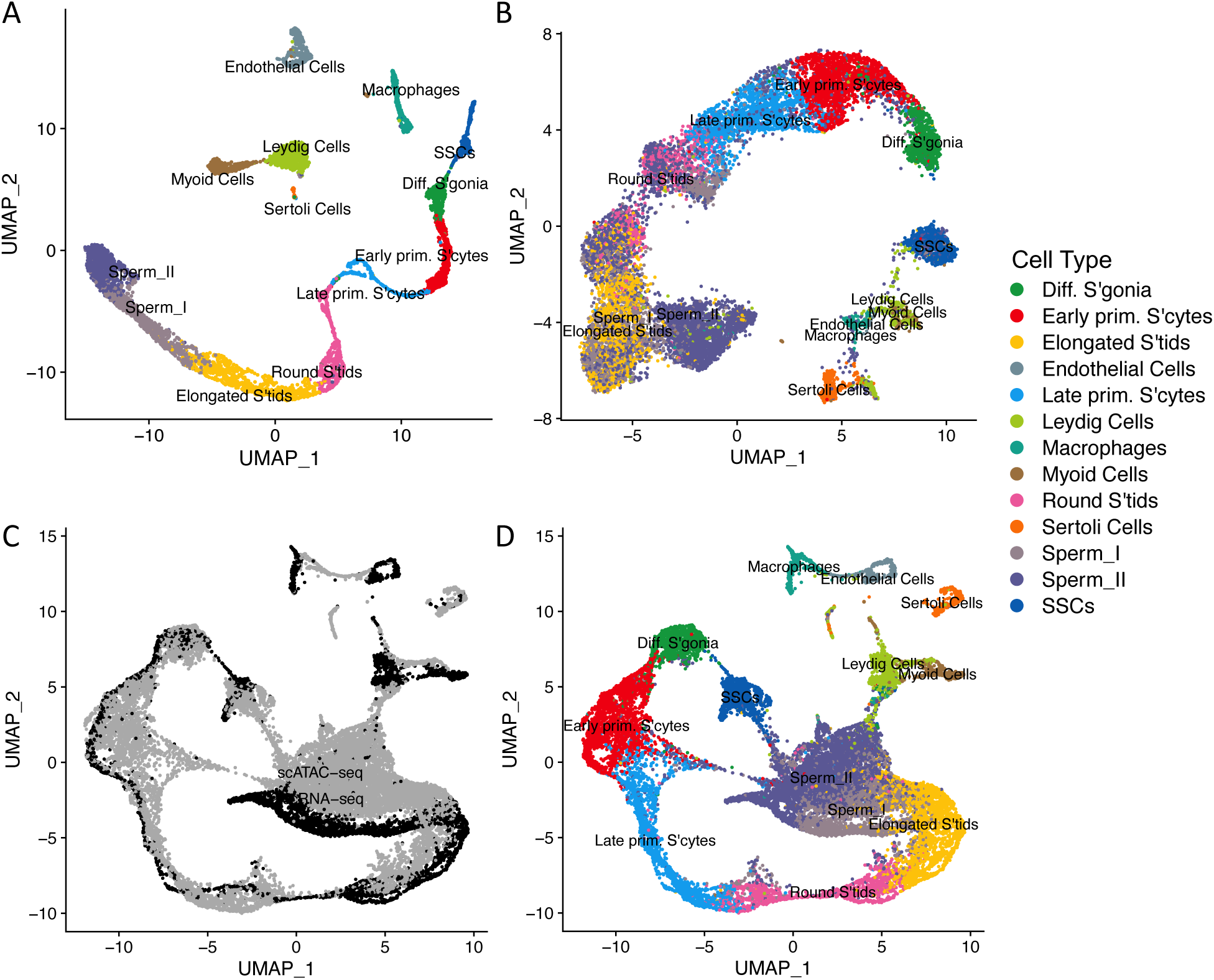
Single cell expression and chromatin accessibility patterns during human spermatogenesis. Dimension reduction (UMAP) plots of scRNA-seq (A) and scATAC-seq (B) dataset; cell state annotations are derived from marker gene expression in (A) (34). The same data are projected onto a single plot, with the population of origin indicated in (C) and cell annotations shown in (D). In (C), scATAC-seq data are shown in grey and scRNA-seq data in black.

In order to infer chromatin loops accurately in any given cell type, it is crucial to assess CTCF binding activity first. CTCF is known to be widely expressed in humans (37, 38), including in spermatogenesis (39). Post-meiotic cells have a highly compacted genome, with histones being replaced by protamines (40) and transcription largely, but not completely, silenced after the histone-to-protamine transition (41). However, CTCF binding is known to be active in the regions of the haploid sperm genome that escape this transition (42). We predicted bound CTCF sites in all cell types based upon established footprinting analysis within ATAC-seq peaks (Methods). The overall trajectories of CTCF activity were similar in the scRNA-seq and scATAC-seq datasets studied here - as expected if CTCF gene expression results in increased binding. Activity was found to be highest in SSCs and before the first meiotic division, including in early primary spermatocytes; it was then dramatically reduced in round spermatids and beyond, in accordance with a general compaction of the genome (Fig. 2A, B). However, CTCF activity could be measured across all cell types, including in mature sperm. In addition, binding site footprinting plots (35) show a shoulder around CTCF-motifs in all cell types, reflecting a higher-than-expected transposase insertion frequency around the CTCF motif also in post-meiotic cells and thus indicating protein binding at the CTCF motif sites, albeit at reduced intensity (Fig. 2C).

**Fig. 2:**
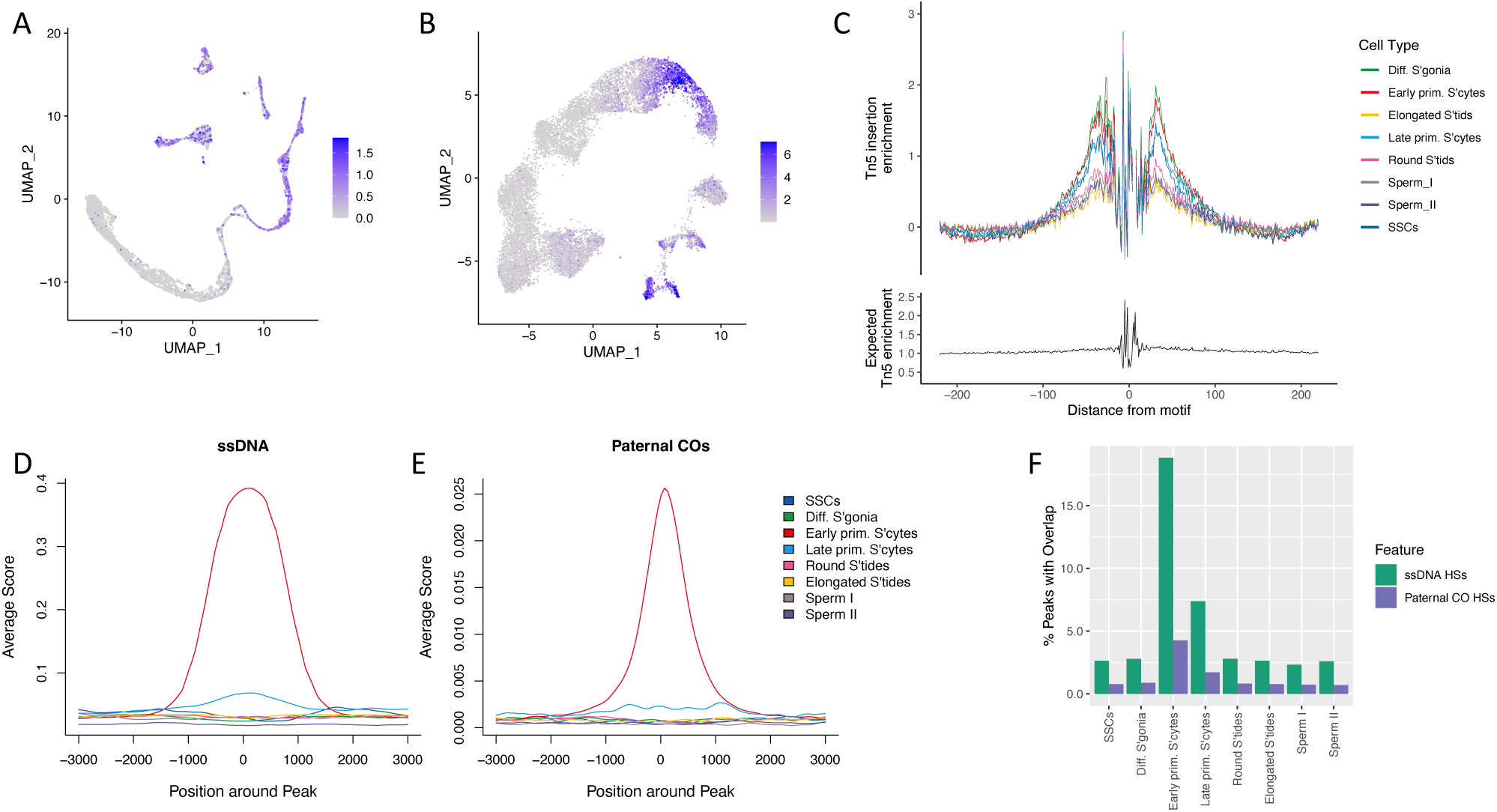
Dynamics of CTCF activity and recombination during meiosis. CTCF activity is visualised in the scRNA-seq (A) and scATAC-seq data (B), using the same embeddings as in Fig. 1. (C) CTCF binding-site footprinting profiles derived from the scATAC-seq data, using Signac (35). Cell-type-specific peaks were profiled for ssDNA overlap (D) as well as paternal cross-over activity (E), using the genomation package in R (43). (F) The 80% reciprocal overlap percentage for all scATAC-seq peaks in a given cell type versus ssDNA HSs (green) and paternal CO hotspots (purple).

Having confirmed CTCF binding activity, we employed a machine learning framework to study the changing chromatin conformation landscape throughout spermatogenesis. We used an established random forest modelling approach for bulk data that allows for *de novo* prediction of chromatin loops (31) and adapted this framework to single cell datasets; we trained the model on aggregated pseudo-bulk scATAC-seq and scRNA-seq data derived from the GM12878 lymphoblastoid cell line (Methods). Using 10-fold cross validation, the model performed well on the GM12878 single cell test data and achieved an area under ROC curve value of 0.95, based on ground truth ChIA-PET and Hi-C datasets of chromatin loops. Loop length and CTCF signal intensities at loop anchors achieved the highest importance scores (Fig. 3).

**Fig. 3:**
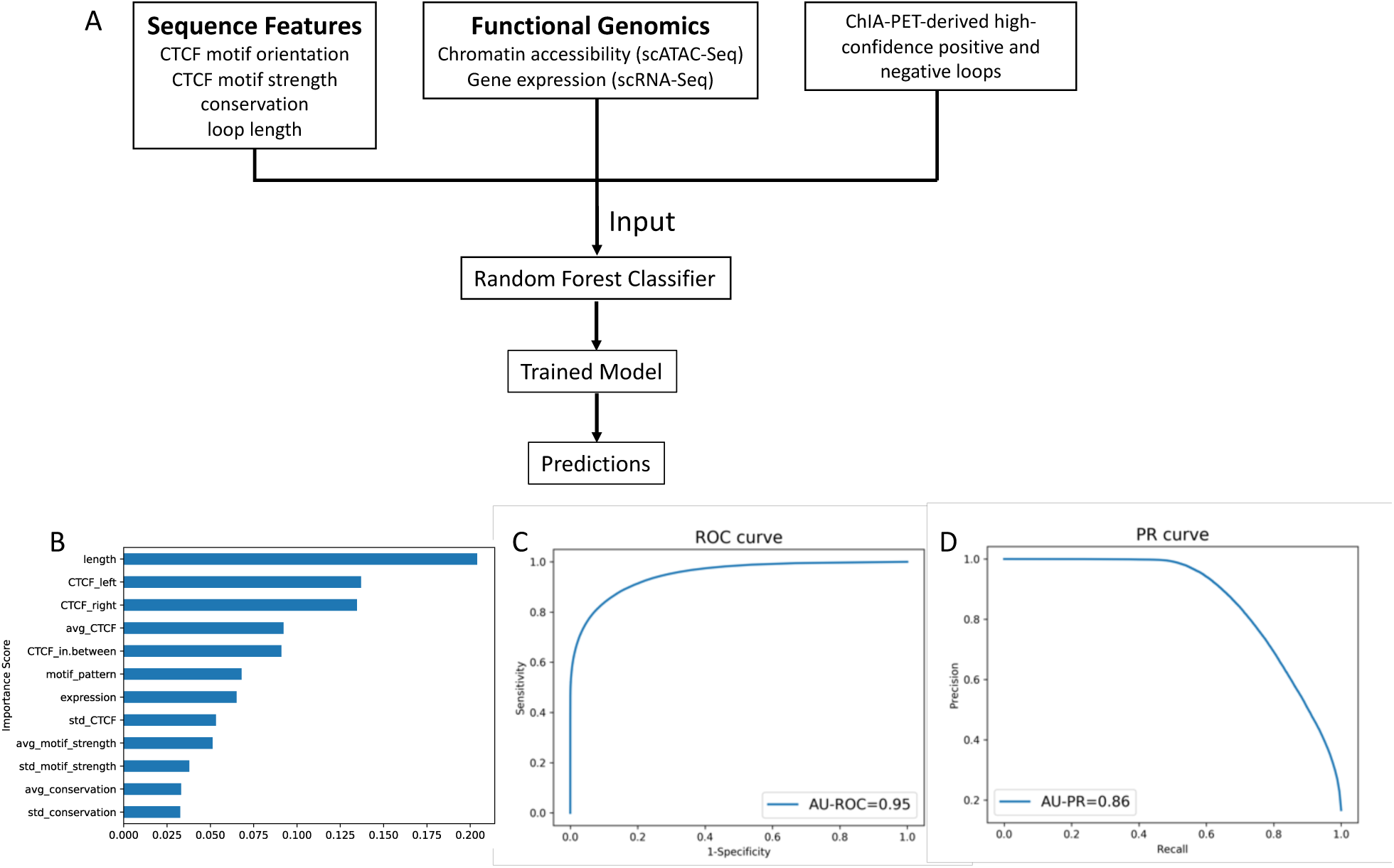
Accurate prediction of chromatin loops using single cell data. A machine learning model was trained on aggregated scRNA-seq and scATAC-seq data in GM12878, using ChIA-PET and Hi-C data as ground truths. (A) Overview of the pipeline, which yields a random forest classifier and can be used to make predictions on a new dataset. (B) Feature importance scores = mean decrease impurity in the training process; “avg” and “std” represent the mean and standard deviation of the signal intensity on both anchors; “left” and “right” represent flanking features and “in-between” is the signal intensity within a loop. (C) Receiver operator characteristic (ROC) curve and (D) Precision recall (PR) curve, using five iterations of 10-fold cross validation.

The model was further validated in an independent single cell dataset from the K562 cell line; using a 99% reciprocal overlap criterion, almost all of the experimentally derived ChIA-PET loops (99%) were predicted by our model (Additional File 3: Table S1). Additional loops that were predicted by the machine learning model but absent in the ChIA-PET data seem to be due to a lack of sequencing depth in the latter, since lowering the minimum PET (paired-end tag) support required for a ChiA-PET loop to be included recovers an increasing number of predicted loops (Additional File 3: Table S1), with a maximum of 59% of predicted loops supported experimentally when ChIA-PET loops are supported by at least 3 PETs.

We further tested the robustness of our model by comparing it to a) a model trained on chromosomal cross-fold validation schemes (44) and b) a model that only used CTCF features for model training (see Methods). Both alternative models resulted in comparable AU-ROC and AU-PR scores as well as a similar estimation of loop numbers in the different spermatogenesis cell types (Additional File 2: Fig. S1).

### Distinct chromatin conformations emerge in preparation for meiosis

We used the trained model to predict chromatin loops in each of the eight cell types separately, using cell-type specific input features of CTCF-binding, chromatin accessibility and gene expression.

Among cell types within the spermatogenesis dataset, the number of predicted loops ranged from 520 in sperm I, to 15,693 in early primary spermatocytes (Fig. 4 and Additional File 1: “Predicted Loops”). However, the number of predicted loops was not a simple consequence of cell numbers in a given cell type. For example, mature sperm (sperm II) had the highest cell count, but, consistent with a compaction of the haploid genome, we found only a moderate number of accessible peaks and a strongly reduced number of predicted loops, relative to other cell types (Fig. 4). Conversely, in the same cells, regions surrounding genes known to be specifically active during the later stages of spermatogenesis showed high signals of accessibility. Examples of such genes are TNP1 and PRM2, both of which are involved in chromatin remodelling of the haploid genome and show accessibility almost exclusively in the haploid phase (Additional File 2: Fig. S2).

**Fig. 4:**
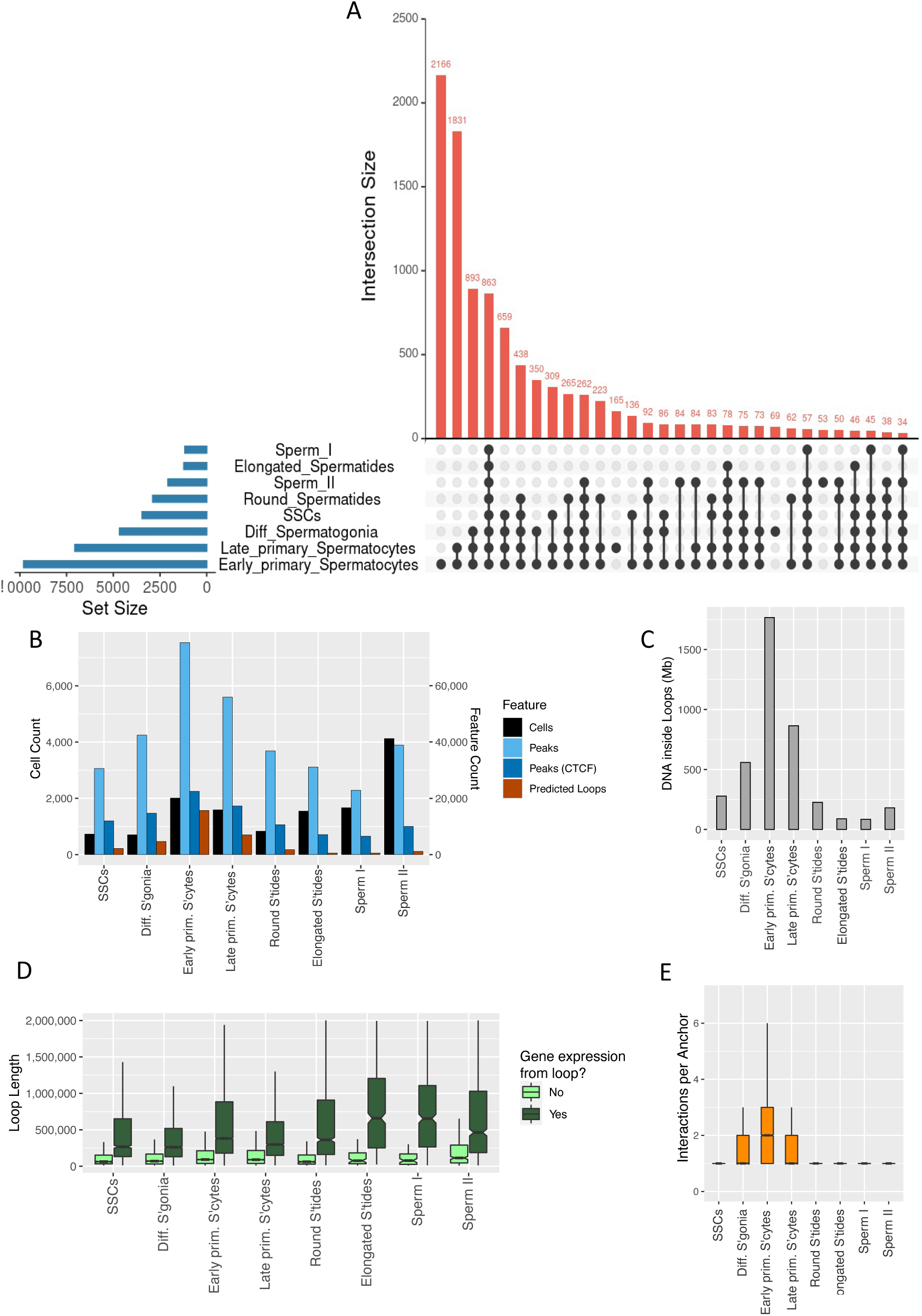
Characteristics of predicted loops in germ cell development. (A) Upset plot of the number of predicted loops in each cell state and their mutual overlap. (B) The total number of cells in the scATAC-seq dataset assigned to a given cell state (black bars, left Y axis). The total number of scATAC-seq peaks called in each of the cell states (light blue), the number of peaks containing CTCF footprints (dark blue) and the number of predicted loops (red) are shown on the right hand Y axis. (C) The total amount of unique DNA covered by loops annotated in each of the germ cell populations. (D) The distribution of loop lengths for loops that contain genes which are expressed, compared to loops that do not contain expressed genes. (E) The number of predicted interacting anchors per CTCF-anchor. Boxplots in (D) and (E) indicate the median (central bar), the 25^th^ and 75^th^ percentiles (boxes), and 95% percentiles (whiskers).

Pairing and synapsis of homologous chromosomes takes place at the early primary spermatocyte stage, and various interconnected aspects of predicted chromatin structure reach notable maxima, including chromatin accessibility (numbers of scATAC-seq peaks), CTCF binding and numbers of chromatin loops (Fig. 4A, B, C). Compared to earlier and later time points, early primary spermatocytes also stood out as having more complex and variable loop structures, with a higher number of interactions measured for a given loop anchor site (Fig. 4E). Chromatin loops in early primary spermatocytes contained by far the largest amount of unique DNA (1,766MB; Fig.

4C), with a median distance of 390Kb between non-overlapping loops. Overall, loops increase in size in the transition from SSCs to early primary spermatocytes - to around half a megabase, on average - and then decrease again in late primary spermatocytes, with the presence of expressed genes being associated with longer loop lengths (Fig. 4D). Note that this association between loop length and expression is not unique to the spermatogenesis dataset, but was also found in the GM12878 training set, with median loop lengths of 87Kb and 476Kb for loops with zero and non- zero expression levels, respectively; Wilcoxon Test: *W* = 4,817,254,898, *p* < 10^^-16^.

### A post-meiotic shift of loops to the telomeres

The chromosomal distribution of CTCF sites and loops differed substantially between cell types: active (i.e. scATAC-seq footprinted) CTCF sites were somewhat biased towards the telomeres at all stages of meiosis, with about 4-5% of CTCF sites falling within the 1% outermost ends of the chromosomes (Fig. 5A). However, in the haploid phase, this bias in the distribution of CTCF is dwarfed by the difference in predicted loop density near the telomeres: in Sperm I, a total of 75% of loops (390 out 520) are predicted to fall into the 1% outermost bins of sequence (Fig. 5B). These loops do not originate post-meiotically but are already present in SSCs (Fig. 4A), suggesting that, in the haploid phase, loops are lost along the length of the chromosomes, rather than specifically gained near the telomeres, resulting in an enrichment of loops near the chromosomal ends. In stark contrast, the most uniform distribution of loops along the chromosomes is found during prophase I of meiosis, in early primary spermatocytes (Fig. 5B, C).

**Fig. 5:**
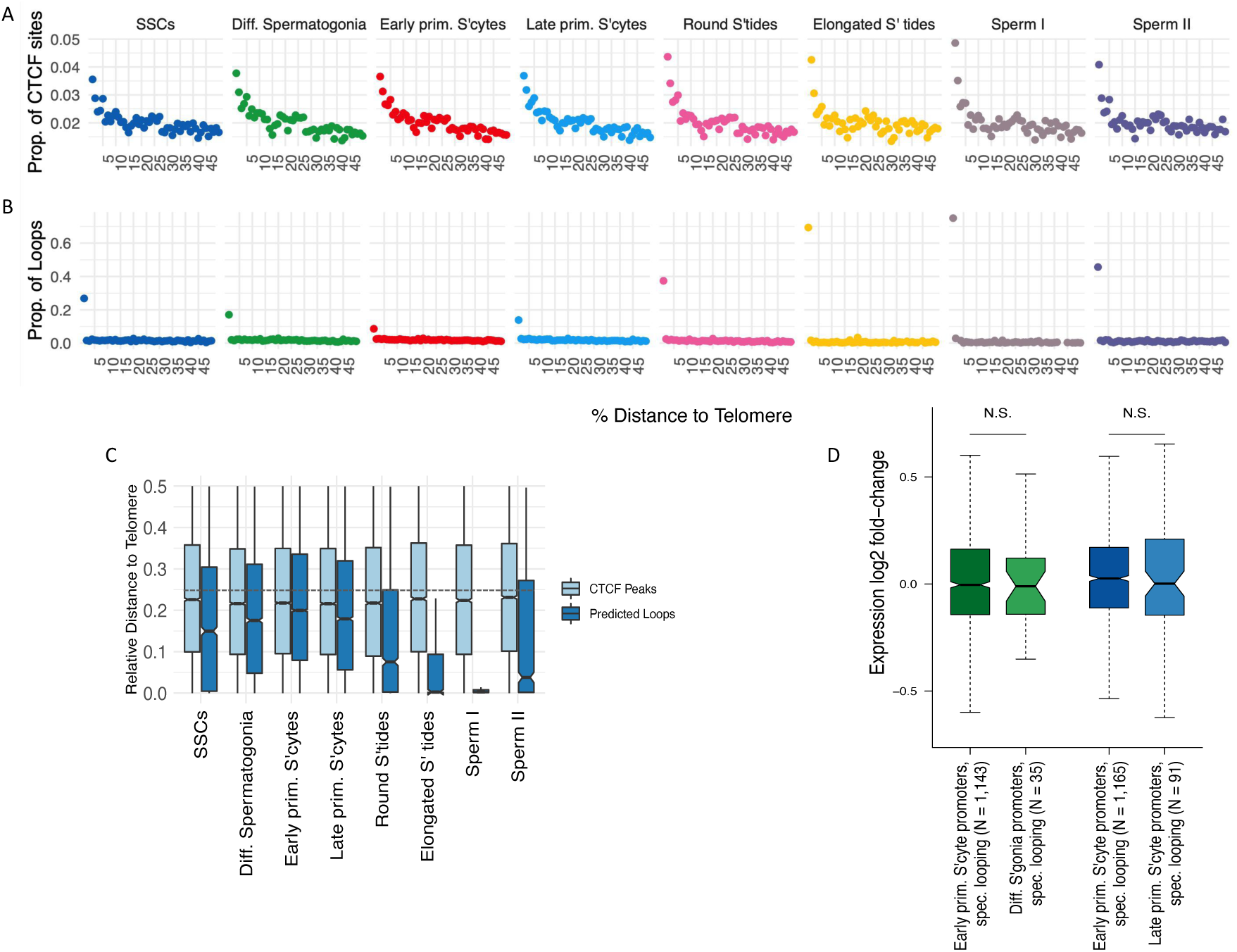
The dynamics of chromatin loop location during spermatogenesis and the impact of looping on gene expression change. In (A and B), chromosomes were divided into 100 even-sized bins; the X axis shows the distance of a given bin to the nearest telomere, relative to chromosome length. On the Y axes, we plot, on a per-cell type basis and across all chromosomes: (A) the proportion of footprinted CTCF sites in a given bin, relative to the total number of footprinted CTCF sites; (B) the proportion of predicted loops in a given bin, relative to the total number predicted loops. (C) The distance to the nearest telomere, relative to chromosome length, is shown for CTCF motifs (light blue) and predicted loops (dark blue). An average relative distance of 0.25 is expected if elements are evenly distributed with respect to telomeric distance, indicated by the dashed line. (D) Looping does not result in gene activation in early primary spermatocytes. Y axis: log2 fold-change of gene expression, with log2 fold-values > 0 indicating higher expression in early primary spermatocytes. The number (N) of promoters located in cell-type specific loops is indicated in each comparison, and the Wilcoxon signed-rank test was used to compare the median log2 fold- change between categories. Green boxplots: early primary spermatocytes versus differentiating spermatogonia; promoters are either in early primary spermatocyte-specific loops (left) or in differentiating spermatogonia-specific loops (right). Blue boxplots: early versus late primary spermatocytes; promoters are either in early (left) or late (right) primary spermatocyte-specific loops.

Overall, these results suggest that chromatin conformation acquires distinct features as CTCF-anchored loops become more abundant in the transition from SSCs to meiotic cells. Loops become more variable between individual cells, reflected by a higher number of interactions per CTCF anchor, increase in size and are more evenly spread across the chromosomes - until they shrink in absolute numbers and are largely confined to the chromosomal ends in haploid cells.

### Chromatin looping does not drive gene expression changes during spermatogenesis

A substantial literature on chromatin loops and related structures, such as TADs (e.g. (45–48), has suggested these structures have roles in gene regulation, coordinating expression changes in functionally related genes by facilitating enhancer-promoter interactions (49–51). We tested this idea by contrasting the gene expression change associated with loops that specifically emerged in the transition “differentiating spermatogonia -> early primary spermatocytes” and “early primary spermatocytes -> late primary spermatocytes”. Further, we only considered genes whose promoters were involved in chromatin loops exclusively in early primary spermatocytes or exclusively in the preceding or subsequent cell type (see upset plot in Fig. 4A). If chromatin looping led to promoter activation, the log2-fold ratio of gene expression would be, on average, larger for genes in early spermatocyte-specific loops. However, this was not the case (Fig. 5D), i.e. loops which emerged specifically in prophase I were not associated with the activation of promoters that reside inside these loops. Similarly, the overlap between a promoter and an emerging loop anchor point (plus minus 1Kb) was not associated with a change in gene expression from differentiating spermatogonia to early primary spermatocytes or from early to late primary spermatocytes (Wilcoxon tests: W = 699, N.S., and W = 231, N.S.).

We further studied the relationship between gene function and chromatin looping by comparing - on a per cell-type level - the gene ontology (GO) enrichments of genes with promoters inside or outside of loops, respectively, ranked by gene expression level (52). GO categories of highly expressed genes turned out to be very similar between the two categories of genes (Additional File 3: Table S2). For example, in sperm I, the top ten enriched GO categories are identical between genes whose promoters are involved in looping versus genes with promoters outside loops; in both cases, gene function of highly expressed genes mainly relates to mRNA processing and translation, reflecting the silencing of most transcription in sperm and a shift towards post-transcriptional processing of existing transcripts (53). Similar results (no difference in GO categories) obtained for the seven other spermatogenesis cell types (Additional File 3: Table S2) and indicates that loops do not contain genes that are functionally different from non-loop genes during meiosis. Another possibility, which could explain changes in looping, is that chromatin conformation dynamics are largely mechanistically associated with aspects of meiosis itself, such as recombination, and we explored this option further.

### Meiotic chromatin accessibility and chromatin looping are major determinants of global recombination patterns

Compared to pre-or post-meiotic germline cell types, ATAC-seq peaks in early primary spermatocytes were most strongly associated with measures of recombination, including ssDNA hotspots (54) as well as paternal cross-over events (55) (Fig. 2D, E and F). A total of 21.6 % of accessible sites (16,244/75,351) in early primary spermatocytes overlapped DMC1-bound ssDNA peaks; genome-wide, the overlap represents a more than 5-fold enrichment compared to random expectation (p = 0.001 with N= 1,000 circular permutations) and shows that double-strand breaks during prophase I of meiosis are often contained within accessible sites of cells that prepare to undergo chromosome pairing (56, 57). In addition, using the MEME de novo search algorithm (58), we retrieved an experimentally derived PRDM9 binding motif as the most significant motif from these DMC1-bound/accessible chromatin sites (E-value = 3.2e-11; Additional File 2: Fig. S3). Next, we investigated how chromatin loops in early primary spermatocytes relate to recombination events. Dividing loops into even-sized bins and aggregating signals across all loops, we find that ssDNA hotspots and paternal cross-over events are enriched at loop anchors, whereas haplotype blocks often end at loop anchors (Fig. 6A, B, C) (59). In an independent analysis and using circular permutations, we quantify the genome-wide enrichment of ssDNA hotspots at loop anchors as 3.6-fold, and the enrichment of paternal cross-over locations as 3.5-fold (Table 1). Conversely, haplotype blocks are depleted at loop anchors relative to genome-wide expectations (Table 1) though loops themselves tend to overlap many haplotype blocks (Fig. 6G), i.e. anchors stand out in their genomic context as disrupting haplotype blocks, indicative of historical recombination events at loop anchor sites.

**Fig. 6:**
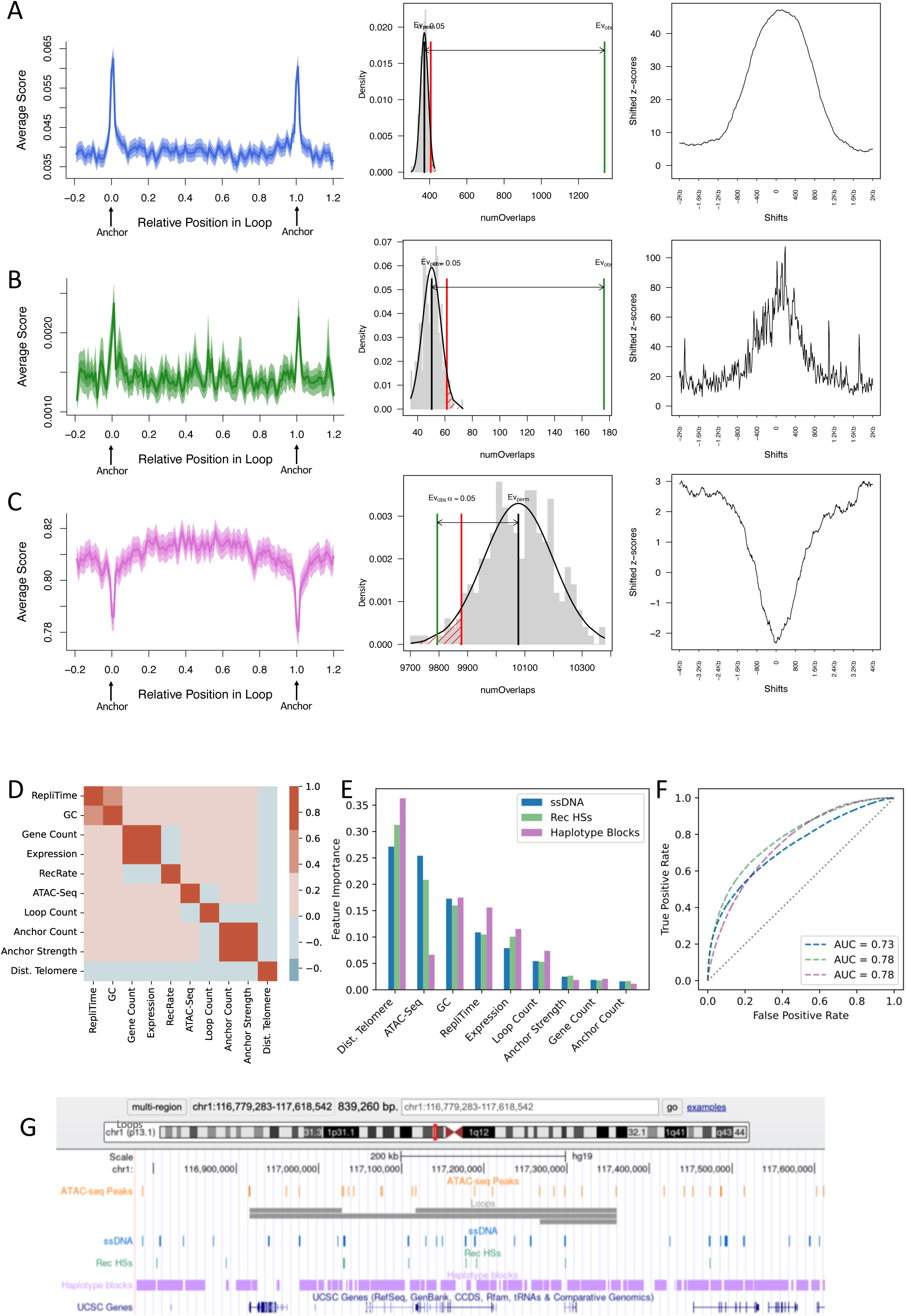
Chromatin looping is associated with DNA breakage and recombination. (A) ssDNA hotspots, (B) paternal crossing over events and (C) East Asian ancestry (EAS) haplotype blocks were intersected with chromatin loops in early primary spermatocytes (from EAS sample donors). In each row, the first plots show average enrichment scores across all loops (plus/minus 20% of loop length), including, as shaded bands, the standard error of the mean and the 95 percent confidence interval for the mean. The second plot in each row is a probability density plot based on 1,000 circular permutations; the observed number of overlaps between loop anchors and (A) ssDNA peaks, (B) paternal crossing over events (C) EAS haplotype blocks is indicated by the green vertical lines; in each case the expected distribution of overlap is shown as a grey histogram. The threshold for statistical significance is indicated by the red vertical lines. The last plot in each row shows the distribution of shifted Z-scores, i.e. the variation in circular permutation Z-scores if loop anchors and ssDNA peaks, paternal crossing over events or EAS haplotype blocks, respectively, were shifted with respect to each other. A distinct peak (or trough) in the local Z-scores indicates that overlaps result from specific local patterns. D) Spearman’s correlation coefficient between genomic features, calculated genome-wide in 1Kb genomic bins. E) Feature Importance Plot for the Random Forest classifier models of ssDNA hotspots (blue), recombination hotspots (green) and EAS haplotype blocks (purple). F) Receiver Operating Characteristic (ROC) curve of all three models. G) Illustrative UCSC genome browser screen shot of the distribution and size of genomic elements. Shown are ATAC-Seq peaks in early primary spermatocytes, predicted loops in early primary spermatocytes, ssDNA hotspots, paternal recombination hotspots, EAS haplotype blocks and UCSC gene models.

**Table 1:**
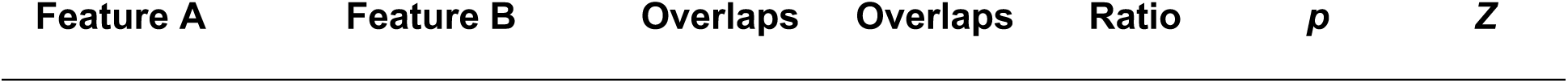
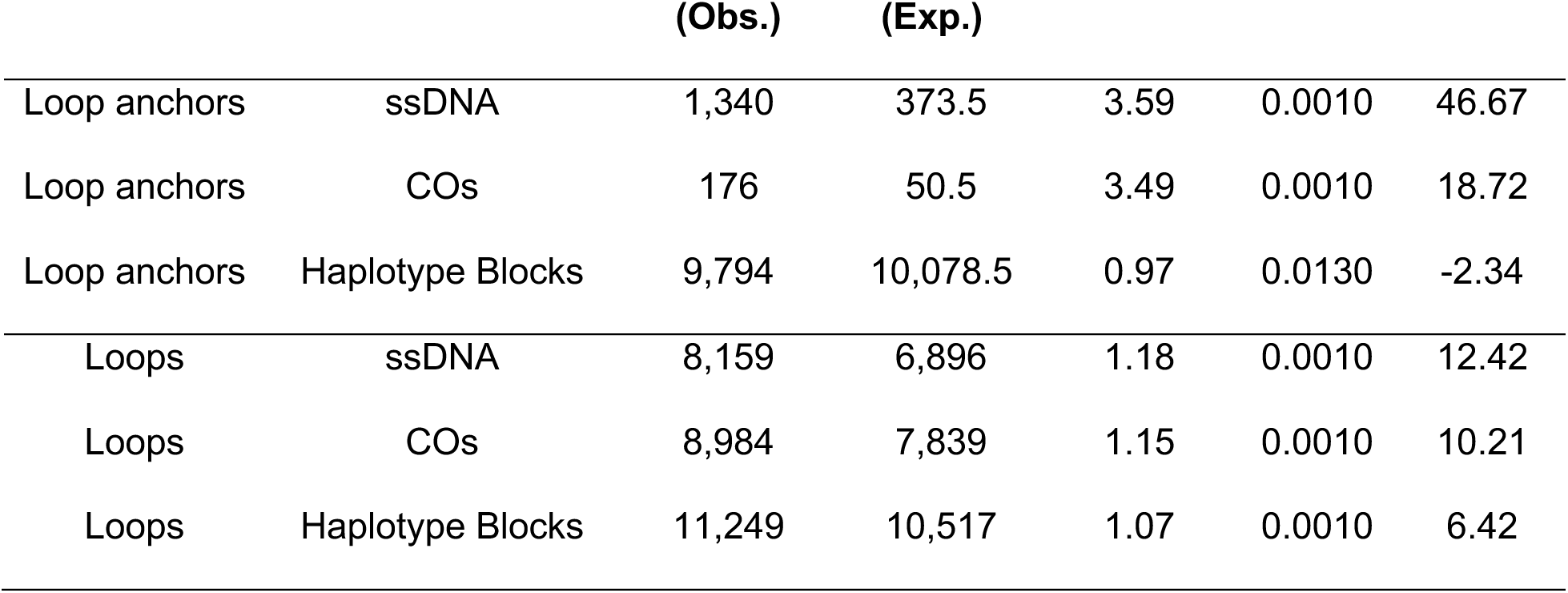
Circular permutations Results. Statistics of overlap between looping and recombination features. The ratio of observed over expected overlaps is shown in column 5, and associated *p-* values and permutation *Z*-scores in columns 6 and 7. All results are based on N = 1,000 permutations.

|  |  | (Obs.) | (Exp.) |  |  |  |
| --- | --- | --- | --- | --- | --- | --- |
| Loop anchors | ssDNA | 1,340 | 373.5 | 3.59 | 0.0010 | 46.67 |
| Loop anchors | COs | 176 | 50.5 | 3.49 | 0.0010 | 18.72 |
| Loop anchors | Haplotype Blocks | 9,794 | 10,078.5 | 0.97 | 0.0130 | -2.34 |
| Loops | ssDNA | 8,159 | 6,896 | 1.18 | 0.0010 | 12.42 |
| Loops | COs | 8,984 | 7,839 | 1.15 | 0.0010 | 10.21 |
| Loops | Haplotype Blocks | 11,249 | 10,517 | 1.07 | 0.0010 | 6.42 |

Although chromatin conformation appears to play a role in recombination, a variety of other features are known to be important in the broad genome-wide patterns of recombination. To investigate the impact of chromatin folding on recombination while controlling for other intercorrelated factors (Fig. 6 D), we employed a set of random forest models to predict genome-wide patterns of recombination in relation to genomic features such as GC content, chromatin accessibility, meiotic replication timing, gene expression levels, chromosomal positioning, as well as variables associated with chromatin loops in prophase I. We modelled, as categorical outcome variables, the presence of a) ssDNA and b) paternal recombination hotspots in 1Kb windows along the genome (based on the data of Pratto et al. (54) and Halldorsson et al. (55)), as well as c) the overlap with EAS haplotype blocks (59). In all three models, the random forest classifier achieved moderate ROC-AUC scores, and telomeric distance was the most important predictor (Fig. 6E, F); this may reflect the fact that paternal cross-over events are biased towards the tip of the chromosomes (60) and, similarly, Subramanian (61) observed an increase in DSB formation in telomere-proximal regions in yeast. Chromatin accessibility ranked highly in the models of ssDNA and recombination hotspots, consistent with the observation that PRDM9 creates nucleosome-depleted regions (56, 57); however, previous studies have lacked accessibility data from meiotic human cells. Local GC content and meiotic replication timing also ranked highly across models, and both features are known to correlate with levels of recombination (62, 63). With respect to looping features, the overlap of a genomic window with the *internal* section of a loop is, in all three models, more important than the number of loops emerging from a given 1kb bin or the predicted anchor strength in a genomic window (Fig. 6). Thus, while the presence of anchors and loops both contain predictive information on recombination, overall, the internal sequence of chromatin loops is more strongly associated with recombination features than the anchor sites that form the boundaries to loops.

It is an open question why some ssDNA sites are selected as sites of crossing-overs, whereas others are repaired without this type of chromosomal exchange (64). Hence, we partitioned ssDNA HSs based on their overlap with paternal recombination HSs and investigated how chromatin looping, chromatin accessibility and nearby gene activity affect the chance for an ssDNA site to be selected as the site of crossover formation. First, we noted that the 16,712 ssDNA HSs that are also recombination HSs (ssDNA+/rec_HSs+) were more likely to overlap chromatin loops in early primary spermatocytes compared to the 7,986 ssDNA HSs that did not act as recombination HSs (ssDNA+/rec_HSs-) (Odds ratio 1.37 (95% confidence intervals [1.32, 1.42]), suggesting a positive association between looping and cross-over formation after double-strand break initiation. Within the set of loop-overlapping ssDNA HSs, associated loop expression levels were *lower* for ssDNA+/rec_HSs+ compared to ssDNA+/rec_HSs- (median loop expression levels of 2.6 and 2.1, respectively; Wilcoxon Test: *W* = 170,923,365, *p* < 10^-16). Similarly, the level of regulatory activity - measured as the number of footprints inside early primary spermatocyte ATAC-seq peaks - was lower at ssDNA+/rec_HSs+ compared to ssDNA+/rec_HSs- sites (a median of 2 *versus* 5 footprints per peak; Wilcoxon Test: W = 90,436,936, p-value < 10^-16^), suggesting that gene expression negatively associates with cross over formation at ssDNA sites. Next, we assessed the enrichment of transcription factors motifs at footprinted ssDNA+/rec_HSs+ versus ssDNA+/rec_HSs- sites (65, 66). The PRDM9 (MA1723.1) motif was enriched at ssDNA+/rec_HSs+ compared to ssDNA+/rec_HSs- sites (OR = 1.14, log10(*p*-value) = 10.24), whereas CTCF (MA0139.1) was enriched among ssDNA+/rec_HSs- sites (OR = 1.35 , log10(*p*-value) = 15.16). In a ranked list analysis (52) across all DNA-binding proteins that were expressed in early primary spermatocytes, functional GO terms associated with "chromatin insulator sequence binding" and "RNA polymerase II proximal promoter sequence-specific DNA binding" were enriched among ssDNA+/rec_HSs- compared to ssDNA+/rec_HSs+ site (Additional File 3: Table S3), again highlighting the negative effect of gene transcription and the presence of insulator sequences on cross over formation. Thus, chromatin looping and gene activity affect downstream processing of double-strand break sites at multiple levels: cross-over formation is favoured *within* loops, but gene activity inside loops counteracts this process, rendering regions with higher regulatory and transcriptional activity less likely to be favoured.

## DISCUSSION

Chromatin loop dynamics during human meiosis are poorly understood and, consequently, the roles of such 3D structures in meiotic cell function are an area of active investigation. Here, we define looping patterns during human spermatogenesis and find dramatic changes between cell types,ranging from expanded loops encompassing much of the genome in early primary spermatocytes (as cytologically described in (5)), to the mature sperm genome, which almost exclusively forms loops near the telomeres In addition, we find that changes in chromatin looping are not strong predictors of gene expression dynamics during meiosis, and no evidence for an enrichment of functional categories of genes inside loops, suggesting that loops largely play an architectural role, primarily facilitating synapsis of homologous chromosomes.

Homologous recombination is invariably linked to the 3D structure of meiotic chromosomes (reviewed in (3)). Here, we use human genomic data to show that early spermatocyte loop structures have an impact on the position of both DNA breakage and cross over events – with DMC1-bound ssDNA sites more likely to result in cross overs if located inside a loop (8). Further, haplotype block ends are enriched near loop ends, reflecting increased linkage disequilibrium (LD) within prophase I loops – in contrast to results of a previous study that investigated chromatin interactions in somatic cells (67). However, we note that haplotype blocks tend to be much smaller than loops, on average (see Fig. 6G), and LD will not extend very far in relation to loop size , hence not be sufficient to maintain favourable combinations of promoter and enhancer variants relevant for prophase I. An enrichment of haplotype block ends near anchor points is, however, consistent with our results that recombination often occurs at anchor sites during meiosis, albeit at lower rates than suggested based on ssDNA sites alone. We have shown that chromatin loop anchor sites can be predicted using a computational approach to exploit previously published single cell expression and chromatin accessibility data. Cell states in the spermatogenesis scATAC-seq dataset were identified by stratifying cells based on their accessibility profiles near marker genes, and footprinting analysis was used to computationally identify CTCF-bound sites – based on prior knowledge of characteristic transposase “footprints” near protein-binding motifs (68). To build even more precise models, it would be desirable to obtain ChIP-seq data of CTCF within individual spermatogenesis cell types in humans, especially since some CTCF-binding sites may be occupied by its meiosis-specific paralogue (BORIS/CTCFL), which does not interact with cohesin (69) but occupies a subset (∼12%) of CTCF motif sites during spermatogenesis (70). In contrast to CTCF sites, however, CTCFL/BORIS-binding sites are not oriented in a convergent orientation and are biased towards more accessible chromatin compared to anchor sites (70) - both of which are factors that form part of our model training.

In our study, computationally determined cell states were further corroborated by the strong association of accessible sites and features of recombination that were only found in early primary spermatocytes – a relationship that is only expected if cell populations are accurately determined. Next, we used the information on CTCF-binding, in combination with other features, to identify chromatin loops in meiotic cell populations. In such pools of cells, our training data showed high accuracy in predicting chromatin looping – despite cohesin (RAD21), which facilitates loop extrusion (71, 72) - not being part of the model. This is consistent with reports that features related to CTCF binding may be sufficient to predict looping in most contexts (32). In addition, looping was accurately predicted in a different cell line with corresponding single cell data.

Meiotic chromatin dynamics are further complicated by the fact that some cohesin subunits are replaced by meiosis-specific molecules (e.g. REC8 instead of RAD21), and the possibility remains that this modification could change the outcome of the loop dynamics that we are aiming to predict. Our model does not take into account large-scale compartment switches and global chromatin changes, which have been observed during mammalian meiosis (20, 21), nor the role of chromosomal axis proteins (73–75) or other factors involved in DSB- initiation, such as the recombination initiation complex (e.g. SPO11), which is known to interact with the chromosomal axis (76, 77). Hence, it would be worthwhile to extend to more multi-faceted machine learning models that also incorporate larger-scale chromatin changes and information on other players that may drive the positioning of recombination events.

Despite these limitations, our model predicts loop expansion in preparation for the first meiotic division, a phenomenon that has also been observed in murine Hi-C data (28). However, despite our model allowing for loops sizes of up to 2MB, we estimate loop sizes in early primary spermatocytes to have a mean length less than 500KB, corroborating the notion that computational methods can detect smaller loops that may be overlooked *in vivo*. Loops are biased towards the tips of the chromosomes already in early primary spermatocytes, where male recombination is known to be enhanced (78). Later during spermatogenesis, in the compacted sperm genome, loops are largely confined to the tips of the chromosomes. This observation is consistent with experimental data which have shown that telomeric regions are the only regions that retain their histones after the histone-to-protamine transition in haploid sperm; these telomeric ends locate at the nuclear membrane and are free to be actively transcribed in sperm (79). In sperm, loop sizes in the compacted part of the genome were often larger in size than loops in prophase I; this is also in line with inactive chromosome regions having, on average, larger loops sizes (8, 18); thus, the phenomenon of enlarged loops in sperm might reflect different processes compared to the expansion of loops in diploid pre-meiotic cells. Overall, our predictions on chromatin looping align with experimental studies (18, 28), while providing the exact positioning of loops in human meiosis - or, rather, a snapshot of loop positions, given that CTCF binds DNA at a much more dynamic way compared to cohesin, with shorter binding times (80), and the process may be more dynamic than captured by our methods Another shortcoming of the data analysed here is that looping of the maternal and paternal chromosomes cannot be distinguished - something that could only be achieved using crosses between inbred lines, such as mouse strains. It would also be beneficial to dissect the different stages that make up prophase I, to paint an even higher resolution picture of chromatin dynamics during meiosis. This would require higher numbers of cells in which to estimate loops in our pseudo-bulk analyses. In this study, the PRDM9 genotype of the three South East Asian tissue donors was unknown – and may be a combination of different alleles, given the high level of polymorphism at the PRDM9 locus; thus, we used the combined set of ssDNA hotspots detected across genotypes (54), which in turn limits the accuracy that any random forest model of ssDNA and recombination hotspots can achieve. Finally, the paternal recombination map was derived from the Icelandic population, which is likely to differ between the recombination map in South East Asians, but genetic maps with a similar level of resolution are not available; again, this sets a limit to the accuracy of our modelling. Nevertheless, the positioning of CTCF sites (and hence that of most loops) is likely to be conserved between European and Asians, and the predicted looping structure should be largely transferable between populations.

## CONCLUSIONS

Computational predictions of DNA folding are a powerful technique to study the process of meiosis and recombination, in cell populations which are otherwise difficult to study. Our predictions of chromatin loops in human meiosis align with experimental data from other organisms, but the genome-wide resolution obtained by our approach is unprecedented and provides novel insights into the processes that drive human diversity and adaptation.

## METHODS

### scRNA-seq processing

Single cell gene expression data of testicular cells were downloaded from the GEO database (Series GSE112013) (34), including the UMI (unique molecular identifier) table for 6,490 individual cells. Cluster identity for each cell was downloaded from (34). Based on marker gene expression, cells had been classified into 13 clusters, including 8 germline cell types at different stages of differentiation as well as 5 somatic cell types (34). Using Seurat 4.2 in R 4.1.3, we created a Seurat object from these data, using the parameters min.cells = 3, min.features = 200 and followed the standard Seurat pipeline including NormalizeData(with normalization.method = "LogNormalize" and scale.factor = 10000), FindVariableFeatures(using selection.method = "vst" and nfeatures = 2000), ScaleData() and RunPCA() , finding variable features (), scaling the data and running PCA, using the variable features. After checking the Elbowplot for the number of dimensions to use, we used FindNeighbors(with dims = 1:20) and RunUMAP(with dims = 1:20). We used the original cell type classification (34) to plot the dimension reduction plot using DimPlot(). Aggregate expression values in each cell type were calculated using Seurat’s AggregateExpression(), which also calls ScaleData(). We used GTFtools_0.9.0 to extract the transcription start sites as defined in the gtf file Homo_sapiens.GRCh38.97.chr.gtf, and the scaled expression data were converted to an input expression Table as required for the machine learning pipeline (https://github.com/ykai16/Lollipop).

### scATAC-seq processing

Raw sequencing and barcode files were downloaded from the SRA sequence archive (SRR21861961, SRR21861962 and SRR21861963) (13) and processed using 10x Genomics Cell Ranger’s cellranger-atac count (version 1.1.0). Using standard filtering of low-quality barcodes, this resulted in 20,296 barcodes being assigned to cells. We loaded the filtered peak barcode matrices from the Cell Ranger output into Seurat 4.2 using Read10X_h5() and performed batch correction using Harmony(with reduction = ’lsi’), followed by RunUMAP(with dims = 2:30 , reduction = ’harmony’), and FindNeighbors(with k.param = 20, prune.SNN = 0.3, n.trees = 30).

Cell states were transferred from the scRNA dataset to the scATAC dataset using Signac 1.10.0 (35). First, we next created a Fragment object with CreateFragmentObject() and calculated GeneActivity() in the scATAC-seq dataset using VariableFeatures() of the scRNA-seq dataset. After normalisation and scaling of the gene activity data using default parameters, we calculated “transfer anchors” between the two datasets, using FindTransferAnchors(with reduction = "cca" and k.filter=NA), i.e. including genes of all lengths. We predicted cell states within the scATAC dataset using TransferData() and added this to the meta data of the scATAC dataset using AddMetaData(). Only cells with a prediction score > 0.4 were kept for further analysis.

UMAP plots including the predicted cell identities was visualised using DimPlot( ), and, for illustrative purposes, we co-embedded the scRNA and scATAC data on the same plot using merge(), followed by ScaleData(), RunPCA() and RunUMAP().

CallPeaks() was used to call ATAC-seq peaks independently in the thirteen predicted cell types. We created a fragment object of cells whose associated cell-type prediction scores were > 0.4, and then created a feature matrix from which we created a new Seurat object, using CreateChromatinAssay() followed by CreateSeuratObject(), excluding blacklisted regions.

We added motif information and footprinting of CTCF using the Seurat Footprint( ) function, with “in.peaks = TRUE” and ran RunChromVAR() to visualise the activity of CTCF in the different cell types using FeaturePlot(). Observed CTCF footprints in the different cell types as well as the expected Tn5 insertion enrichment was plotted using the PlotFootprint() function. FindTopFeatures() was used to obtain a measure of peak activity, calculated as the percentile of activity in each cell type separately.

FilterCells() was used to obtain separate bed files of all fragments in the different cell types separately, and fragments overlapping CTCF motif-containing peaks were extracted using bedtools intersect, yielding a file of sequence fragments overlapping CTCF-binding sites.

All output from the initial scRNA-seq and scATAC-se analysis was lifted to the hg19 assembly using liftOver tool.

### GM12878 single cell data processing

We processed single cell data of the GM12878 cell line in a way analogous to the testis datasets.

scATAC-seq data came from the 10X Genomics website, a 1:1 mixture of 1:1 Mixture of Human GM12878 and Mouse EL4 Cells (https://www.10xgenomics.com/resources/datasets/1-1-mixture-of-fresh-frozen-human-gm12878-and-mouse-el4-cells-2-standard) with the associated data summary available at https://cf.10xgenomics.com/samples/cell-atac/2.1.0/10k_hgmm_ATACv2_nextgem_Chromium_X/10k_hgmm_ATACv2_nextgem_Chromium_X_web_summary.html. First, we extracted the 4,595 cells annotated as “human” (as opposed to “mouse”) and created a chromatin assay, then a Seurat object from those human-derived data. We ran FindTopFeatures() and created a fragment object based on human-only fragments. Using AddMotifs() and Footprint(), we obtained all estimated CTCF-binding sites in GM12878, including, as a measure of peak strength, the percentile of cut sites. A separate bed file contained all sequence fragments overlapping those CTCF peaks.

We investigated the correspondence between our estimated CTCF-binding sites versus experimentally derived CTCF-binding sites in GM12878 by making use of encode ChIP-seq data (accession numbers ENCFF797SDL, ENCFF951PEM, ENCFF796WRU and ENCFF827JRI) and confirmed that the majority of sites were shared between the different datasets (Additional File 2: Fig. S4). Notably, the 12,148 sites that were shared among all 4 ChIP-seq experiments but not found in the scATAC-seq data had lower signal values compared to the 20,066 ChIP-seq-scATAC-seq shared sites (median signal values of 102.4 and 201.5, respectively; Wilcoxon test: W = 193,625,454, p < 2.2e-16 ). Conversely, sites that were only present as CTCF footprinted peaks had a somewhat reduced peak strength compared to sites that were shared with the ChIP-seq data (median percentile cut sites of 0.49 and 0.53, respectively; W = 115,733,156, p = 1.703e-09).

Single cell RNA for GM12878 was downloaded from the GEO database (dataset GSM3596321) (81). We processed the data in the same way as the testis-derived data, including creating a Seurat object from the gene expression matrix (min.cells = 3, min.features = 200), running NormalizeData(), FindVariableFeatures(selection.method = "vst", nfeatures = 2000), ScaleData() and RunPCA(). We then ran AggregateExpression() to obtain scaled gene expression values for the 18,170 genes provided in the gene expression matrix.

### Adapting a machine learning pipeline to single cell data

To train the random forest classifier (31), we used ChIA-PET (Geo dataset GSM1872886) and Hi-C data (Geo dataset GSE63525) from the GM12878 cell line.

First, we trained the model on 90,211 “true” ChiA-Pet supported loops with a median size of 164,546bp, using the provided scripts “prepare_training_interactions.py”, “add_features.py” and “train_model.py” (available at https://github.com/ykai16/Lollipop). The parameters of the model were relaxed so as to allow loops of up to 2MB, by setting “maxLength" in “prepare_training_interactions.py” to 2000000. Positive and negative training interactions were characterised by training features, which included scaled gene expression values, scATAC-seq peaks as well as footprinted CTCF sites and their relative peak strength. Additional features included the CTCF motif score (sequence similarity to the consensus motif, calculated by the original authors using Fimo (82)) as well as the phastCon motif conservation of the CTCF motifs (http://hgdownload.cse.ucsc.edu/goldenpath/hg19/phastCons100way) (83).

We further modified the pipeline to encompass single cell data: the bed file associated with the CTCF peaks contained all fragments overlapping peaks (rather than raw reads overlapping peaks), and, accordingly, we adjusted the “shift” parameter in “add_features.py” to half the median size of the sequence fragments (=136/2), and also removed the strand information associated with raw reads files. Further, since we have fewer input variables compared to the published pipeline, we changed the parameters “max_depth” and “max_features” in the RandomForestClassifier() function of “train_models.py” to the default values of “None” and "sqrt", respectively (Scikit-learn package (84)).

The model was evaluated computationally using 10-fold cross validation with the ‘StratifiedKFold’ strategy to ensure that each fold had the same proportion of class labels as the original dataset (84). Cross-validation results, ROC and PR curves were automatically created.

Last, using the script “make_denovo_predictions.py” (also available at https://github.com/ykai16/Lollipop), the trained classifier was used make *de novo* predictions of chromatin looping in the eight spermatogenesis cell types separately (SSCs, Differentiating Spermatogonia, Early Primary Spermatocytes, Late Primary Spermatocytes, Round Spermatides, Elongated Spermatides, Sperm I and Sperm II), including the same set of input features as in the model trained on GM12878 – again, allowing for interactions of up to 2Mb by setting “distance_distal = 2000000” and adjusting the “shift” parameter in “lib.py” to half the median size of the sequence fragments (136/2).

### Testing the robustness of the machine learning model

We modified the “train_models.py” script further and performed chromosome-based cross-validations instead of random sub-sampling, which can sometimes lead to over-fitting (44). Using the GroupKFold method from sklearn.model_selection with n_splits=5, we set group_kfold.split(X, y, groups=chroms) to split the data based on chromosomal location, thereby preventing data leakage from the training into the test sets.

Similarly, we also trained a more simplified model (based on the original re-sampling scheme (31)) by removing gene expression and ATAC-seq peaks from the set of training features – and keeping only CTCF peak strength, conservation and motif orientation as input features. Both alternative models were evaluated using ROC and PR curves.

### Validation of predicted loops in an independent single cell dataset

We tested the performance of the modified random forest classifier on the K562 cell line, using published scATAC-seq (85), scRNA-seq (86) and ChIA-PET high confidence CTCF-mediated loop data (87) (geo datasets GSE162690, GSM5687481, GSM5374829). A Seurat object was created from the scRNA-seq data using the provided barcodes, features and matrix data, and aggregate gene expression values were calculated in the same way as for the GM12878 and spermatogenesis datasets above. Raw fastq files were extracted from the scATAC-seq data, i.e. the “high loading” mixture of ten human cell lines, using Cell Ranger’s bamtofastq, and then processed with cellranger-atac count (version 2.1.0). The 2,066 barcodes associated with the K562 cell line (85) were used to subset this dataset, followed by peak calling and CTCF foot-printing analysis in Seurat and Signac. Chromatin loops were predicted in the K562 cell line based on the random forest classifier trained on the single cell GM12878 data, i.e. in a fashion analogous to the spermatogenesis dataset but using K562- specific input parameters of chromatin accessibility (276,933 peaks), CTCF-binding (70,127 active sites), and gene expression. The resulting predicted chromatin loops were compared to “ground truth” ChIA-PET loops of the K562 cell line; first, we calculated the midpoints of each anchor region in the ChIA-PET dataset and defined each experimental loop as extending between these two midpoints. We only included loops of 10KB to 2MB in size, as these limits are also set in the machine learning model. To assess the impact of sequencing depth in the ChIA-PET data, we varied the minimum number of PETs required to call a ChIA-PET loop. Then, we used bedtools intersect to count the number of predicted loops that were also reported experimentally, using a 99% reciprocal overlap criterion.

### Enrichment Calculations

Meiotic double-strand break maps (ssDNA maps) in human testes were obtained from (54). Paternal cross-over maps came from (55). Based on paternal recombination rate estimates (55), recombination hotspots were defined as regions with a recombination rate >= 9.85 centiMorgan per Mb (>= 10x the genome-wide average).

To calculate haplotype blocks, we downloaded the phase 3 1000 Genomes SNP and Indel file from http://hgdownload.cse.ucsc.edu/gbdb/hg19/1000Genomes/phase3/. Plink version 1.90b4 was used to filter the vcf files for bi-allelic variants called in the East Asian Ancestry (EAS) population, and haplotype blocks were calculated using the --blocks command with no-pheno-req no-small-max-span --blocks-max-kb 2000. Haplotype blocks of a minimum size of 1kb were used for further analysis.

The R package genomation 1.26.0 (43) was used to visualize enrichments of sequence features near accessible sites and predicted loops in early primary spermatocytes, including the overlap between loop features and ssDNA overlap, paternal cross-over frequency and haplotype block ends. In the case of accessible sites, we used the resize() function with “width = 6000” and “fix = "center"”), followed by the ScoreMatrix() function to calculate genomic enrichments around the accessible sites in all eight cell types. For loops in early primary spermatocytes, we considered the full length of loop coordinates plus/minus 20% of the loop length, and then divided each such region into 140 even-sized bins. The ScoreMatrixBin() function was used to calculate genomic enrichments for each of the 120 bins – which varied in absolute size among loops.

The R package regioneR 1.26.1 (88) was used to perform circular permutations of sets of genomic regions, i.e., early primary spermatocyte loop anchor points or loops, respectively, *versus* haplotype blocks, ssDNA peaks and paternal cross over events. We used the permTest() function with parameters ntimes = 1000, randomize.function = circularRandomizeRegions, evaluate.function = numOverlaps, genome = hg38_masked, alternative = “auto,” where hg19_masked=getBSgenome(”BSgenome.Hsapiens.UCSC.hg19.masked”). This yielded a genome-wide estimate of enrichment of overlap between features as measured by Z-scores as well as the ratio of observed to expected overlaps.

### Motif Analysis

To facilitate *de novo* motif discovery, the “rgt-hint footprinting” algorithm was applied to scATAC-seq peaks in early primary spermatocytes and their corresponding bam file of aligned reads. The thus obtained 481,840 footprints were intersected with ssDNA peaks (54), and FASTA sequences of the 93,681 footprints inside ssDNA peaks were subjected to the MEME *de novo* motif search algorithm (58), using the parameters -nmotifs 10 -minw 6 -maxw 50. Motifs were passed onto the Tomtom algorithm (89), using the default parameters -min- overlap 5 -mi 1 -dist pearson -evalue -thresh 10.0, thus searching for similar, previously described, motifs among the *de novo* motifs inside ssDNA HSs.

Footprints inside ssDNA peaks were further processed using the JASPAR enrichment tool (https://jaspar.elixir.no/enrichment/) (65, 66), searching against a known database of DNA binding motifs, i.e. all available human JASPAR 2022 TFBS sets (available at https://zenodo.org/records/6860555) with detectable expression levels in early primary spermatocytes. Footprints were divided into those that overlapped either ssDNA+/rec_HSs- or ssDNA+/rec_HSs+ sites. Next, we used the JASPAR_enrich.sh script with the “twoSets” parameter, testing for differential TFBS enrichment between the two sets of genomic regions. Motifs were ranked by their odds ratio of enrichment in ssDNA+/rec_HSs+ *versus* ssDNA+/rec_HSs- and *vice versa*, and the ranked lists of corresponding transcription factors were compared using the GOrilla tool at https://cbl-gorilla.cs.technion.ac.il (52).

### Calculation of Telomere Distances

To calculate the distance of a genomic feature (CTCF motif of loops) to the nearest telomere, we calculated the distance of the 5’ and 3’ end of the feature to the 5’ and 3’ end of the chromosome, and selected the smallest of these four distances. To calculate the *relative* distance to the nearest telomere with respect to chromosome length, we simply divided telomere distance by chromosome length. Thus, if features are randomly distributed, we expect, on average, a relative telomeric distance of 0.25.

### GO analysis with respect to looping features

GO analysis was carried out using the GOrilla tool at https://cbl-gorilla.cs.technion.ac.il (52). For all eight germline cell types, we extracted all genes whose promoter sequence overlapped a predicted chromatin loop in the given cell type, and ranked these genes by their scaled gene expression values. A second list of genes included all genes whose promoter sequence did *not* overlap a predicted chromatin loop in a given tissue, and this list was also ranked by gene expression levels. Next, we ran the GOrilla tool on each ranked list of genes.

### Gene expression and looping changes

We examined if changes chromatin loops in early primary spermatocytes were associated with gene expression changes by considering cell types with “private” loops, i.e. loops that were predicted to be present only in a single cell type; this included 2,166 loops in early primary spermatocytes, 165 loops in late primary spermatocytes, 69 in differentiating spermatogonia and 53 in sperm II (Fig. 4).

We applied Seurat’s FindMarker() function to calculate gene expression changes (log2 fold-change of expression) in the comparison of early primary spermatocytes *versus* each of the other three cell types, using the parameters “min.diff.pct = 0, logfc.threshold = 0, min.pct = 0.1, min.cells.group = 3”. Next, we intersected gene promoters (defined as the transcription start site plus/minus 2kb) with loops that were found a) exclusively in early primary spermatocytes and b) exclusively in differentiating spermatogonia, late primary spermatocytes or sperm II, respectively. The median log2 fold-change of expression was compared between the two categories of genes - promoters in early primary spermatocyte-specific loops or promoters in the respective other cell type’s loops - using a Wilcoxon sign rank test.

### Random Forest Model

Random forests for classification and regression were implemented using the scikit-learn python package (84). First, the hg19 genome was sub-divided into 2,599,489 1KB windows, excludeding encode blacklisted regions as well as windows containing “NA” values for any of the predictor variables. For each window, the following sequence features were calculated (using bedtools map, betools intersect and custom R and python scripts, respectively): the average GC content; the distance to the nearest telomere; the average meiotic replication time derived from meiotic S-phase nuclei (DNA content: 2-4C; SYCP3-positive and DMRT1- negative) (63); the number of genes overlapping the window (based on the ensemble annotation “Homo_sapiens.GRCh37.82.chr.gff3”); the number of predicted loops and loop anchor points overlapping the genomic window; the maximum strength of CTCF-anchors in the interval; scaled gene expression values of genes and the average scATAC-seq signal within a window. Cell-type specific features (ATAC-seq signal, gene expression values and looping features) were specific to early primary spermatocytes. Loop anchors were defined as 5kb blocks of sequence surrounding the anchor CTCF site, and loop anchors occurring “internal” of other loops (i.e. overlapping larger loops), were excised from the genomic interval of a loop, in order to avoid the presence of anchors within loops; loops were pruned by 2.5kb on either end, to remove upstream and downstream anchors. Spearman correlation coefficients between numerical variables of the model were visualised using the seaborn package (90).

In the random forest classifier model of “ssDNA hotspot overlap”, we modelled the presence of double-strand break hotspots from (54), and included all hotspots observed in any of the AA1, AA2, AB1, AB2 and AC individuals (with the genotype referring to the PRDM9 allele of the respective donor).

In the random forest classifier model of “paternal recombination hotspot overlap”, we modelled the presence of hotspots defined as genomic intervals for which the paternal recombination rate was >= 9.85 cM/Mb (55).

In both models, the input data were subdivided into training and test sets (70% of data in the training set) and best model hyperparameters were searched using the RandomizedSearchCV() function of the scikit-learn package, sampling 10 hyperparameter values of “n_estimators” (values between 50 and 200) and “max_depth” ( values between 1 and 20), with 5-fold cross-validation. Following this procedure, we built final “best” models with 111 trees and a maximum depth of 18 for ssDNA hotsposts, 62 trees with a maximum depth of 16 for recombination hotspots, and 158 trees with a maximum depth of 19 for haplotype blocks.

## SUPPLEMENTARY INFORMATION

### Additional file 1

*Genomic coordinates (hg19) of predicted chromatin loops in the eight germline cell types*.

### Additional file 2

Figure S1. A comparison of machine learning models to predict CTCF loops.

*Figure S2.* DNA accessibility signal around germ cell marker genes.**Figure S3.** Motif matches within footprints of early primary spermatocyte peaks.

*Figure S4.* Comparison of CTCF ChIP-seq peaks versus footprinted scATAC-seq peaks in GM12878.

### Additional file 3

***Table S1.*** *Confidence of loop predictions using ChIA-PET as a control*.

***Table S2.*** *GO enrichments in eight germline cell types*.

***Table S3.*** *Functional GO enrichments at ssDNA sites that do not progress as recombination hotspots (ssDNA+/rec_HS- compared to ssDNA+/rec_HS+)*.

## DECLARATIONS

### Ethics approval and consent to participate

Not applicable.

## Consent for publication

Not applicable.

## Availability of data and materials

All data used in this study are publicly available, from sources indicated in the manuscript.

## Authors’ contributions

The study was conceived and data were analyzed by V.B.K.; V.B.K. and C.A.S. wrote the manuscript.

## Acknowledgements

Not applicable.

## References

1. Felsenstein J. The evolutionary advantage of recombination. Genetics. 1974;78(2):737–56.

2. Fisher RA. The Genetical Theory of Natural Selection. Oxford: Clarendon Press; 1930.

3. Zickler D, Kleckner N. Meiosis: Dances Between Homologs. Annu Rev Genet. 2023;57:1–63.

4. Hunter N. Meiotic Recombination: The Essence of Heredity. Cold Spring Harb Perspect Biol. 2015;7(12).

5. Zickler D, Kleckner N. Meiotic chromosomes: integrating structure and function. Annu Rev Genet. 1999;33:603–754.

6. Hinch AG, Becker PW, Li T, Moralli D, Zhang G, Bycroft C, et al. The Configuration of RPA, RAD51, and DMC1 Binding in Meiosis Reveals the Nature of Critical Recombination Intermediates. Mol Cell. 2020;79(4):689–701 e10.

7. Baudat F, de Massy B. Regulating double-stranded DNA break repair towards crossover or non-crossover during mammalian meiosis. Chromosome Res. 2007;15(5):565–77.

8. Jin X, Fudenberg G, Pollard KS. Genome-wide variability in recombination activity is associated with meiotic chromatin organization. Genome Res. 2021;31(9):1561–72.

9. Myers S, Bottolo L, Freeman C, McVean G, Donnelly P. A fine-scale map of recombination rates and hotspots across the human genome. Science. 2005;310(5746):321- 4.

10. Jeffreys AJ, Kauppi L, Neumann R. Intensely punctate meiotic recombination in the class II region of the major histocompatibility complex. Nat Genet. 2001;29(2):217–22.

11. Paigen K, Petkov P. Mammalian recombination hot spots: properties, control and evolution. Nat Rev Genet. 2010;11(3):221–33.

12. Kaiser VB, Semple CA. Chromatin loop anchors are associated with genome instability in cancer and recombination hotspots in the germline. Genome Biol. 2018;19(1):101.

13. Wu X, Lu M, Yun D, Gao S, Chen S, Hu L, et al. Single-cell ATAC-Seq reveals cell type-specific transcriptional regulation and unique chromatin accessibility in human spermatogenesis. Hum Mol Genet. 2022;31(3):321–33.

14. Lieberman-Aiden E, van Berkum NL, Williams L, Imakaev M, Ragoczy T, Telling A, et al. Comprehensive mapping of long-range interactions reveals folding principles of the human genome. Science. 2009;326(5950):289-93.

15. Fullwood MJ, Liu MH, Pan YF, Liu J, Xu H, Mohamed YB, et al. An oestrogen-receptor-alpha-bound human chromatin interactome. Nature. 2009;462(7269):58-64.

16. Li Y, Haarhuis JHI, Sedeno Cacciatore A, Oldenkamp R, van Ruiten MS, Willems L, et al. The structural basis for cohesin-CTCF-anchored loops. Nature. 2020;578(7795):472-6.

17. Dixon JR, Jung I, Selvaraj S, Shen Y, Antosiewicz-Bourget JE, Lee AY, et al. Chromatin architecture reorganization during stem cell differentiation. Nature. 2015;518(7539):331-6.

18. Zuo W, Chen GM, Gao ZM, Li S, Chen YY, Huang CH, et al. Stage-resolved Hi-C analyses reveal meiotic chromosome organizational features influencing homolog alignment. Nat Commun. 2021;12(1).

19. Hogarth CA, Evanoff R, Mitchell D, Kent T, Small C, Amory JK, et al. Turning a Spermatogenic Wave into a Tsunami: Synchronizing Murine Spermatogenesis Using WIN 18,446. Biol Reprod. 2013;88(2).

20. Wang Y, Wang H, Zhang Y, Du Z, Si W, Fan S, et al. Reprogramming of Meiotic Chromatin Architecture during Spermatogenesis. Mol Cell. 2019;73(3):547–61 e6.

21. Alavattam KG, Maezawa S, Sakashita A, Khoury H, Barski A, Kaplan N, et al. Attenuated chromatin compartmentalization in meiosis and its maturation in sperm development. Nat Struct Mol Biol. 2019;26(3):175–84.

22. Vara C, Paytuvi-Gallart A, Cuartero Y, Le Dily F, Garcia F, Salva-Castro J, et al. Three-Dimensional Genomic Structure and Cohesin Occupancy Correlate with Transcriptional Activity during Spermatogenesis. Cell Rep. 2019;28(2):352–67 e9.

23. Pugacheva EM, Kubo N, Loukinov D, Tajmul M, Kang S, Kovalchuk AL, et al. CTCF mediates chromatin looping via N-terminal domain-dependent cohesin retention. Proc Natl Acad Sci U S A. 2020;117(4):2020–31.

24. Xiao T, Wallace J, Felsenfeld G. Specific sites in the C terminus of CTCF interact with the SA2 subunit of the cohesin complex and are required for cohesin-dependent insulation activity. Mol Cell Biol. 2011;31(11):2174–83.

25. Xu H, Beasley M, Verschoor S, Inselman A, Handel MA, McKay MJ. A new role for the mitotic RAD21/SCC1 cohesin in meiotic chromosome cohesion and segregation in the mouse. EMBO Rep. 2004;5(4):378–84.

26. Cheng G, Pratto F, Brick K, Li X, Alleva B, Huang M, et al. High resolution maps of chromatin reorganization through mouse meiosis reveal novel features of the 3D meiotic structure. bioRxiv. 2024.

27. Grey C, de Massy B. Chromosome Organization in Early Meiotic Prophase. Front Cell Dev Biol. 2021;9:688878.

28. Patel L, Kang R, Rosenberg SC, Qiu Y, Raviram R, Chee S, et al. Dynamic reorganization of the genome shapes the recombination landscape in meiotic prophase. Nat Struct Mol Biol. 2019;26(3):164–74.

29. Fudenberg G, Imakaev M, Lu C, Goloborodko A, Abdennur N, Mirny LA. Formation of Chromosomal Domains by Loop Extrusion. Cell Rep. 2016;15(9):2038–49.

30. Matthews BJ, Waxman DJ. Computational prediction of CTCF/cohesin-based intra-TAD loops that insulate chromatin contacts and gene expression in mouse liver. Elife. 2018;7.

31. Kai Y, Andricovich J, Zeng Z, Zhu J, Tzatsos A, Peng W. Predicting CTCF-mediated chromatin interactions by integrating genomic and epigenomic features. Nat Commun. 2018;9(1):4221.

32. Xi W, Beer MA. Loop competition and extrusion model predicts CTCF interaction specificity. Nat Commun. 2021;12(1):1046.

33. Xu H, Yi X, Fan X, Wu C, Wang W, Chu X, et al. Inferring CTCF-binding patterns and anchored loops across human tissues and cell types. Patterns (N Y). 2023;4(8):100798.

34. Guo J, Grow EJ, Mlcochova H, Maher GJ, Lindskog C, Nie X, et al. The adult human testis transcriptional cell atlas. Cell Res. 2018;28(12):1141–57.

35. Stuart T, Srivastava A, Madad S, Lareau CA, Satija R. Single-cell chromatin state analysis with Signac. Nat Methods. 2021;18(11):1333–41.

36. Korsunsky I, Millard N, Fan J, Slowikowski K, Zhang F, Wei K, et al. Fast, sensitive and accurate integration of single-cell data with Harmony. Nat Methods. 2019;16(12):1289–96.

37. Filippova GN, Lindblom A, Meincke LJ, Klenova EM, Neiman PE, Collins SJ, et al. A widely expressed transcription factor with multiple DNA sequence specificity, CTCF, is localized at chromosome segment 16q22.1 within one of the smallest regions of overlap for common deletions in breast and prostate cancers. Genes Chromosomes Cancer. 1998;22(1):26–36.

38. Phillips JE, Corces VG. CTCF: master weaver of the genome. Cell. 2009;137(7):1194–211.

39. Hernandez-Hernandez A, Lilienthal I, Fukuda N, Galjart N, Hoog C. CTCF contributes in a critical way to spermatogenesis and male fertility. Sci Rep. 2016;6:28355.

40. Moritz L, Hammoud SS. The Art of Packaging the Sperm Genome: Molecular and Structural Basis of the Histone-To-Protamine Exchange. Front Endocrinol (Lausanne). 2022;13:895502.

41. Dadoune JP, Siffroi JP, Alfonsi MF. Transcription in haploid male germ cells. Int Rev Cytol. 2004;237:1–56.

42. Torres-Flores U, Hernandez-Hernandez A. The Interplay Between Replacement and Retention of Histones in the Sperm Genome. Front Genet. 2020;11:780.

43. Akalin A, Franke V, Vlahovicek K, Mason CE, Schubeler D. Genomation: a toolkit to summarize, annotate and visualize genomic intervals. Bioinformatics. 2015;31(7):1127–9.

44. Xi W, Beer MA. Local epigenomic state cannot discriminate interacting and non-interacting enhancer-promoter pairs with high accuracy. PLoS Comput Biol. 2018;14(12):e1006625.

45. Hansen AS, Cattoglio C, Darzacq X, Tjian R. Recent evidence that TADs and chromatin loops are dynamic structures. Nucleus. 2018;9(1):20–32.

46. Rao SS, Huntley MH, Durand NC, Stamenova EK, Bochkov ID, Robinson JT, et al. A 3D map of the human genome at kilobase resolution reveals principles of chromatin looping. Cell. 2014;159(7):1665–80.

47. Kaiser VB, Semple CA. When TADs go bad: chromatin structure and nuclear organisation in human disease. F1000Res. 2017;6.

48. Dixon JR, Selvaraj S, Yue F, Kim A, Li Y, Shen Y, et al. Topological domains in mammalian genomes identified by analysis of chromatin interactions. Nature. 2012;485(7398):376–80.

49. Dixon JR, Gorkin DU, Ren B. Chromatin Domains: The Unit of Chromosome Organization. Mol Cell. 2016;62(5):668–80.

50. Symmons O, Uslu VV, Tsujimura T, Ruf S, Nassari S, Schwarzer W, et al. Functional and topological characteristics of mammalian regulatory domains. Genome Res. 2014;24(3):390–400.

51. Popay TM, Dixon JR. Coming full circle: On the origin and evolution of the looping model for enhancer-promoter communication. J Biol Chem. 2022;298(8):102117.

52. Eden E, Navon R, Steinfeld I, Lipson D, Yakhini Z. GOrilla: a tool for discovery and visualization of enriched GO terms in ranked gene lists. BMC Bioinformatics. 2009;10:48.

53. Ren X, Chen X, Wang Z, Wang D. Is transcription in sperm stationary or dynamic? J Reprod Dev. 2017;63(5):439–43.

54. Pratto F, Brick K, Khil P, Smagulova F, Petukhova GV, Camerini-Otero RD. DNA recombination. Recombination initiation maps of individual human genomes. Science. 2014;346(6211):1256442.

55. Halldorsson BV, Palsson G, Stefansson OA, Jonsson H, Hardarson MT, Eggertsson HP, et al. Characterizing mutagenic effects of recombination through a sequence-level genetic map. Science. 2019;363(6425).

56. Baker CL, Walker M, Kajita S, Petkov PM, Paigen K. PRDM9 binding organizes hotspot nucleosomes and limits Holliday junction migration. Genome Res. 2014;24(5):724–32.

57. Mansisidor AR, Risca VI. Chromatin accessibility: methods, mechanisms, and biological insights. Nucleus. 2022;13(1):236–76.

58. Bailey TL, Johnson J, Grant CE, Noble WS. The MEME Suite. Nucleic Acids Res. 2015;43(W1):W39–49.

59. Genomes Project C, Auton A, Brooks LD, Durbin RM, Garrison EP, Kang HM, et al. A global reference for human genetic variation. Nature. 2015;526(7571):68-74.

60. Broman KW, Murray JC, Sheffield VC, White RL, Weber JL. Comprehensive human genetic maps: individual and sex-specific variation in recombination. Am J Hum Genet. 1998;63(3):861–9.

61. Subramanian VV, Zhu X, Markowitz TE, Vale-Silva LA, San-Segundo PA, Hollingsworth NM, et al. Persistent DNA-break potential near telomeres increases initiation of meiotic recombination on short chromosomes. Nat Commun. 2019;10(1):970.

62. Eyre-Walker A. Recombination and mammalian genome evolution. Proc Biol Sci. 1993;252(1335):237-43.

63. Pratto F, Brick K, Cheng G, Lam KG, Cloutier JM, Dahiya D, et al. Meiotic recombination mirrors patterns of germline replication in mice and humans. Cell. 2021;184(16):4251–67 e20.

64. Pazhayam NM, Turcotte CA, Sekelsky J. Meiotic Crossover Patterning. Front Cell Dev Biol. 2021;9:681123.

65. Castro-Mondragon JA, Riudavets-Puig R, Rauluseviciute I, Lemma RB, Turchi L, Blanc-Mathieu R, et al. JASPAR 2022: the 9th release of the open-access database of transcription factor binding profiles. Nucleic Acids Res. 2022;50(D1):D165-D73.

66. Sheffield NC, Bock C. LOLA: enrichment analysis for genomic region sets and regulatory elements in R and Bioconductor. Bioinformatics. 2016;32(4):587–9.

67. Whalen S, Pollard KS. Most chromatin interactions are not in linkage disequilibrium. Genome Res. 2019;29(3):334–43.

68. Buenrostro JD, Giresi PG, Zaba LC, Chang HY, Greenleaf WJ. Transposition of native chromatin for fast and sensitive epigenomic profiling of open chromatin, DNA-binding proteins and nucleosome position. Nat Methods. 2013;10(12):1213–8.

69. Pugacheva EM, Rivero-Hinojosa S, Espinoza CA, Mendez-Catala CF, Kang S, Suzuki T, et al. Comparative analyses of CTCF and BORIS occupancies uncover two distinct classes of CTCF binding genomic regions. Genome Biol. 2015;16(1):161.

70. Sleutels F, Soochit W, Bartkuhn M, Heath H, Dienstbach S, Bergmaier P, et al. The male germ cell gene regulator CTCFL is functionally different from CTCF and binds CTCF-like consensus sites in a nucleosome composition-dependent manner. Epigenetics Chromatin. 2012;5(1):8.

71. Kim Y, Shi Z, Zhang H, Finkelstein IJ, Yu H. Human cohesin compacts DNA by loop extrusion. Science. 2019;366(6471):1345-9.

72. Davidson IF, Bauer B, Goetz D, Tang W, Wutz G, Peters JM. DNA loop extrusion by human cohesin. Science. 2019;366(6471):1338-45.

73. Biot M, Toth A, Brun C, Guichard L, de Massy B, Grey C. Principles of chromosome organization for meiotic recombination. Mol Cell. 2024;84(10):1826–41 e5.

74. Pelttari J, Hoja MR, Yuan L, Liu JG, Brundell E, Moens P, et al. A meiotic chromosomal core consisting of cohesin complex proteins recruits DNA recombination proteins and promotes synapsis in the absence of an axial element in mammalian meiotic cells. Mol Cell Biol. 2001;21(16):5667–77.

75. Zickler D, Kleckner N. Recombination, Pairing, and Synapsis of Homologs during Meiosis. Cold Spring Harb Perspect Biol. 2015;7(6).

76. Panizza S, Mendoza MA, Berlinger M, Huang L, Nicolas A, Shirahige K, et al. Spo11- accessory proteins link double-strand break sites to the chromosome axis in early meiotic recombination. Cell. 2011;146(3):372–83.

77. Yadav VK, Claeys Bouuaert C. Mechanism and Control of Meiotic DNA Double-Strand Break Formation in S. cerevisiae. Front Cell Dev Biol. 2021;9:642737.

78. Bherer C, Campbell CL, Auton A. Refined genetic maps reveal sexual dimorphism in human meiotic recombination at multiple scales. Nat Commun. 2017;8:14994.

79. Zalenskaya IA, Bradbury EM, Zalensky AO. Chromatin structure of telomere domain in human sperm. Biochem Biophys Res Commun. 2000;279(1):213–8.

80. Hansen AS, Pustova I, Cattoglio C, Tjian R, Darzacq X. CTCF and cohesin regulate chromatin loop stability with distinct dynamics. Elife. 2017;6.

81. Osorio D, Yu X, Yu P, Serpedin E, Cai JJ. Single-cell RNA sequencing of a European and an African lymphoblastoid cell line. Sci Data. 2019;6(1):112.

82. Grant CE, Bailey TL, Noble WS. FIMO: scanning for occurrences of a given motif. Bioinformatics. 2011;27(7):1017–8.

83. Rosenbloom KR, Armstrong J, Barber GP, Casper J, Clawson H, Diekhans M, et al. The UCSC Genome Browser database: 2015 update. Nucleic Acids Res. 2015;43(Database issue):D670-81.

84. Pedregosa F, Varoquaux G, Gramfort A, Michel V, Thirion B, Grisel O, et al. Scikit-learn: Machine Learning in Python. J Mach Learn Res. 2011;12:2825–30.

85. Granja JM, Corces MR, Pierce SE, Bagdatli ST, Choudhry H, Chang HY, et al. ArchR is a scalable software package for integrative single-cell chromatin accessibility analysis. Nat Genet. 2021;53(3):403–11.

86. Zhao S, Hong CKY, Myers CA, Granas DM, White MA, Corbo JC, et al. A single-cell massively parallel reporter assay detects cell-type-specific gene regulation. Nat Genet. 2023;55(2):346–54.

87. Tang Z, Luo OJ, Li X, Zheng M, Zhu JJ, Szalaj P, et al. CTCF-Mediated Human 3D Genome Architecture Reveals Chromatin Topology for Transcription. Cell. 2015;163(7):1611–27.

88. Gel B, Diez-Villanueva A, Serra E, Buschbeck M, Peinado MA, Malinverni R. regioneR: an R/Bioconductor package for the association analysis of genomic regions based on permutation tests. Bioinformatics. 2016;32(2):289–91.

89. Gupta S, Stamatoyannopoulos JA, Bailey TL, Noble WS. Quantifying similarity between motifs. Genome Biology. 2007;8(2):R24.

90. Waskom ML. seaborn: statistical data visualization. Journal of Open Source Software. 2021;6(60):3021.

