## Additional File 2 for "CTCF-anchored chromatin loop dynamics during human meiosis"

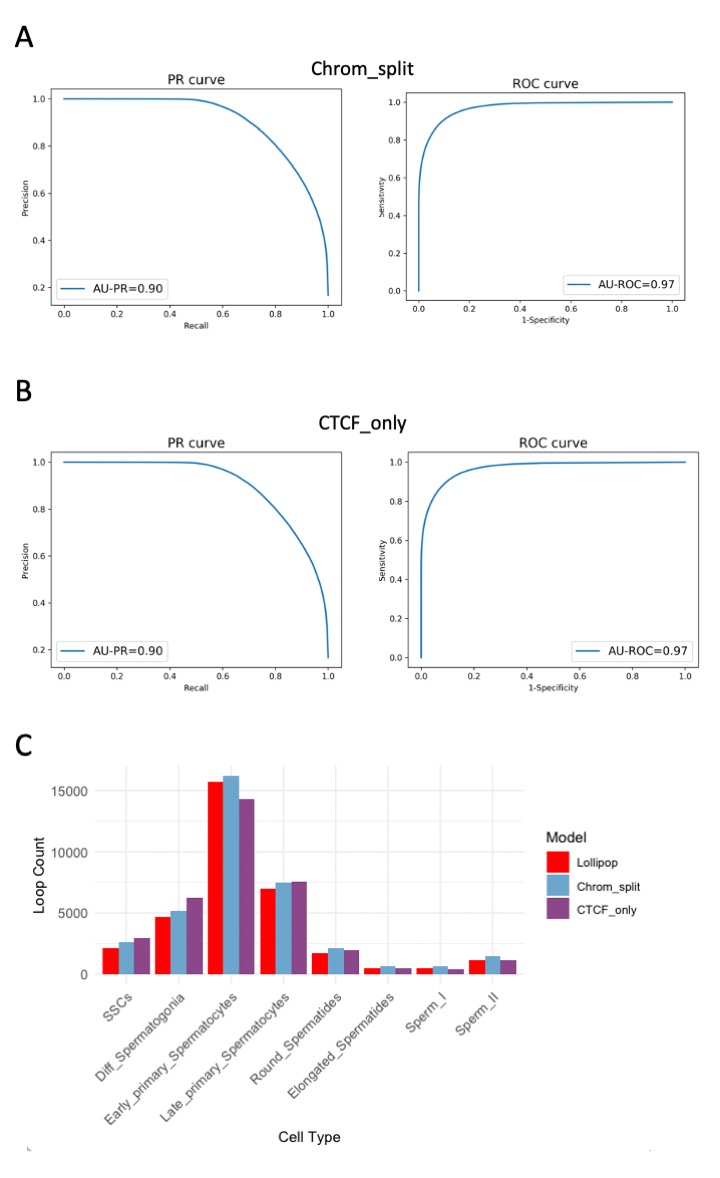

**Supplementary Figure S1: A comparison of machine learning models to predict CTCF loops.**

Model performances (PR and ROC curves) of models trained (A) with chromosomal training/test splits – the “Chrom_split model” - and (B) a reduced model that only uses CTCF-related features as training features – the “CTCF_only model”. (C) Comparing predictions of the number of CTCF-anchored loops based on (A) and (B) to those of the main model used in the manuscript (the “Lollipop” model).

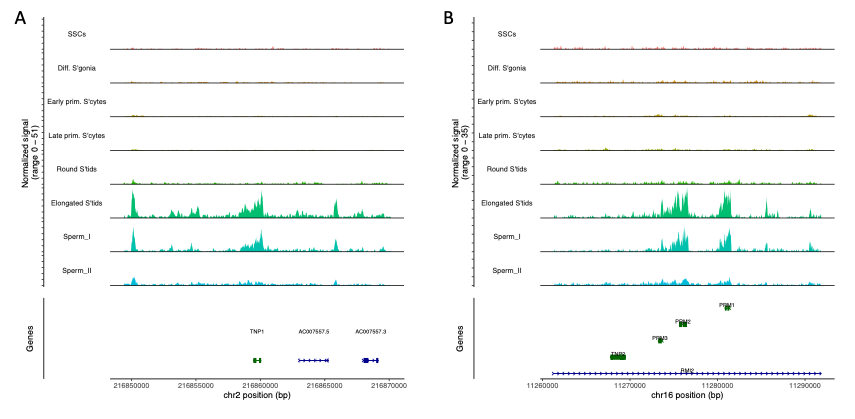

**Supplementary Figure S2: DNA accessibility signal around germ cell marker genes.** Accessibility signals are shown for each of the eight germline cell types separately, along the sequence su­­­rrounding the TNP1 (A) and PRM2 (B) gene. Both genes are preferentially expressed post-meiotically (34) and involved in the histone to protamine replacement during the haploid phase of spermatogenesis.

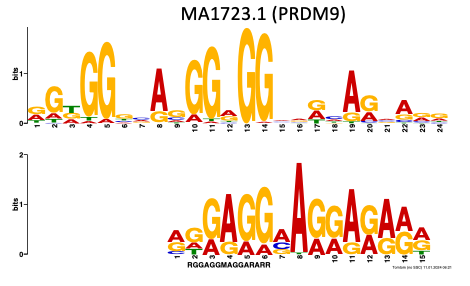

**Supplementary Figure S3:** **Motif matches within footprints of early primary spermatocyte peaks.** The PRDM9 motif in the JASPAR 2022 CORE database (top) and its matching motif inside footprints that also overlap ssDNA hotspots (bottom). Tomtom similarity q-value = 0.019, Benjamini-Hochberg corrected.

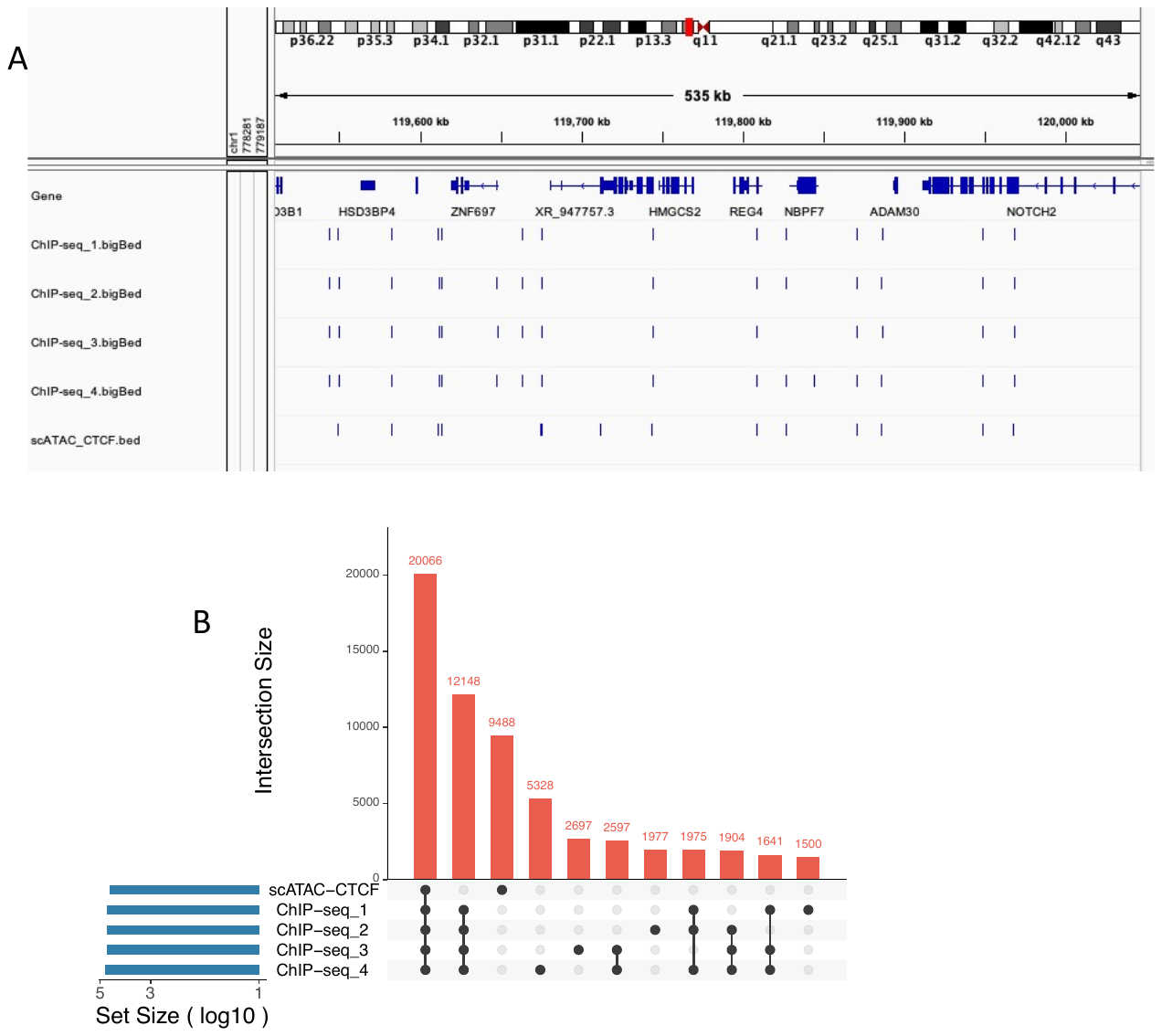

**Supplementary Figure S4: Comparison of CTCF ChIP-seq peaks *versus* footprinted scATAC-seq peaks in GM12878. (**A) An IGV browser screenshot shows the locations of GM12878 ChIP-seq peaks as well as scATAC-seq peaks containing a footprinted CTCF motif. (B) An upset plot of the same data shows the intersection sizes between the datasets. The encode ChIP-seq datasets used in this comparison include datasets ENCFF797SDL (ChIP-seq_1), ENCFF951PEM (ChIP-seq_2), ENCFF796WRU (ChIP-seq_3) and ENCFF827JRI (ChIP-seq_4).
